# BioGAIP: A Scalable, User-Friendly and Robust LLM-Powered Multi-Agent System for Automated Bioinformatics Tasks

**DOI:** 10.64898/2026.05.16.720484

**Authors:** Jiayu Zhang, Pengfei Guo, Guanghui Jiang, Mengyu Zhou, Gang Wei, Ting Ni

## Abstract

The rapid explosion of large-scale, high-throughput biological data has created an urgent demand for efficient analysis pipelines. Traditional bioinformatics approaches, while powerful, often require specialized computational expertise, placing them out of reach for bench biologists. Large Language Models (LLMs) offer new possibilities for automating complex reasoning and tool integration, yet existing LLM-based solutions have not sufficiently lowered this barrier, and expert-level analysis remains inaccessible to most nonexperts. Here, we present BioGAIP, an LLM-powered agent that integrates expert-level reasoning within an end-to-end platform for bioinformatics tasks. By coupling optimized autonomous agents with full graphical interfaces, BioGAIP transforms complex analytical workflows into an automated, user-friendly, and low-intervention process with natural language input. Key features of BioGAIP include dynamic information retrieval, automatic environment configuration, and self-directed design of analysis pipelines, making large-scale multi-omics analysis highly accessible. Built on agent-based client-server architecture, BioGAIP ensures secure resource management and supports heavy computational demands. Extensive evaluations on diverse published datasets demonstrate that BioGAIP reliably recapitulates established biological insights and shows strong potential for novel discovery. By democratizing complex bioinformatics workflows, BioGAIP accelerates accessible data-driven discovery for both experts and nonexperts.

## Induction

The biology field is increasingly inundated with massive volumes of data generated by diverse high-throughput multi-omics technologies (e.g., genomics, proteomics, and metabolomics), which are rapidly expanding in both size and complexity to zettabyte scales^1^, thereby creating an urgent need for efficient processing and analysis^2,3^. Traditional bioinformatics approaches, while powerful, often require sophisticated computational skills, posing significant challenges to bench biologists and researchers without extensive bioinformatics training. The expansion of bioinformatics has rendered it a resource-intensive discipline that characterized by significant resource demands, necessitating robust computational infrastructure and advanced technical expertise. Managing such extensive datasets not only requires establishing specialized computing environments, intricate analytical pipelines, and fine-tuned parameters, but also dealing the additional complexity of integrating diverse data types in multi-omics studies. Traditionally, addressing these multifaceted challenges has heavily relied on the labor-intensive efforts of bioinformatics experts, demands profound cross-disciplinary knowledge to bridge biology, computer science, and statistics. While community-driven initiatives such as nf-core^4^, Galaxy^5^, and OMICtools^6^ have made considerable progress in standardizing and simplifying analysis workflows, the technical barrier remains prohibitively high for wet-lab biologists and clinicians.

A breakthrough came in 2017, with the introduction of the transformer deep neural network architecture was introduced, which fundamentally revolutionized the extraction of long-range contextual relationships in textual data^7^. This innovation laid the foundation for large language models (LLMs)^7^. As a cutting-edge artificial intelligence (AI) method, LLMs have demonstrated capabilities that extend far beyond natural language processing to include robust reasoning, extensive knowledge integration, and skilled code generation. Following the release of ChatGPT-3.5 in 2022, LLMs were initially adopted as conversational chatbots and interactive "copilots" for researchers^7,8^. However, this copilot paradigm inherently limits the broader potential of LLMs. In this mode, LLMs encounter difficulties in complex multi-step workflows that involve non-linguistic computation and ultimately lack the capacity for end-to-end automation. Subsequent advances in prompt engineering and autonomous agent architectures have mitigated this limitations, facilitating the transition from passive chatbot models to proactive agents^9,10^. An LLM-based agent is an autonomous system that utilize language models as its central cognitive engine, leverage techniques such as prompt engineering, environmental awareness, and knowledge integration to iteratively plan tasks, solve problems, invoke external tools, and dynamically interact with other resources^9^. By enabling automated, high-performance analytical pipelines across diverse scientific disciplines, these agent technologies have emerged as a primary driver of LLM-assisted scientific discovery. Consequently, there is solid recognition and excitement regarding the transformative potential of AI for Science (AI4Science)^8^. Several such agents have been specifically developed for bioinformatics, including DrBioRight 2.0^11^ for cancer proteomics, CellAgent^12^ for single cell RNA sequencing (scRNA-seq) and spatial transcriptomics, and BioInformatics Agent (BIA) ^13^ for general bioinformatics assistance. Beyond biology, systems like SciToolAgent^14^ have been deployed across diverse scientific disciplines to automate complex workflows using knowledge graphs and predefined tools.

Despite the significant potential of LLM agents in bioinformatics, their current applications remain limited. Most existing tools suffer from multiple limitations. First, most are confined to specific applications, (e.g., scRNA-seq), relying on predefined toolkits or lack systematic validation on complex multi-omics data processing. Second, these systems continue to impose substantial operational barriers, requiring intricate configuration and specialized expertise, while offering limited capacity for dynamic reasoning and multi-step inference^15^. These shortcomings are exacerbated by critical concerns regarding model reliability, transparency, and security, including hallucinations, unaccountable outcomes, and risks associated with executing code in sensitive computational environments^9,16,17^. Such deficiencies may prevent existing solutions from effectively meeting real-world research demands, which increasingly require the integration of diverse data types and cross-disciplinary insights. Moreover, to maximize broad accessibility, LLM agent-based bioinformatics must present a minimal technical barrier, ideally functioning as out-of-the-box and end-to-end solutions. Achieving this demands advanced planning, contextual troubleshooting, domain-aware inference, intuitive user interfaces, stringent security architectures, and robust cross-platform compatibility. These capabilities remain largely out of reach for current agents.

To overcome the limitations of existing LLM agents and meet the growing demand for accessible, automated, and secure bioinformatics analysis, we introduce BioGAIP (**Bio**informatics **G**enerative **AI P**latform). BioGAIP is a cross-platform, out-of-the-box, interactive container-based system engineered for LLM agent deployment. Guided by the core principles of user-**f**riendliness, **e**lasticity, **s**ecurity, and **s**calability (FESS), BioGAIP comprises three decoupled components: BioAG (**bio**informatics **ag**ents), BioLauncher (a graphical setup wizard) and BioWorker (an execution proxy) (Fig. 1a-b). The BioAG component integrates a graphical interactive user interface (BioAG UI), and a backend agents system built upon the AutoGen^18^ framework (Fig. 2, detail in Methods). Through its integrated agent manager module, BioAG reads configurations for both pre-defined and user-customized agents as well as LLMs client configurations. It facilitates user interaction via an intuitive web-based graphic interface allowing users to seamlessly submit tasks, monitor real-time progress, and intervene as a human-in-the-loop to guide the agentic workflow. Compared to existing tools, BioGAIP i demonstrates superiority across multiple technical dimensions (Table 1), including comprehensive graphical interfaces, support for a broad spectrum of analytical tasks, optimized ensemble of built-in agents, an isolated execution proxy, and robust customizability (detailed in the Methods). These advancements enable fully automated, end-to-end bioinformatics analysis with minimal human intervention. Evidence from our evaluations indicates that the system has the capacity to automate complex tasks with a single or a few user prompts. Collectively, these design features and capabilities endow BioGAIP has with the potential to lower technical barriers, enable the seamless analysis of complex omics data and significantly accelerate researches across diverse fields.

**Fig. 1:**
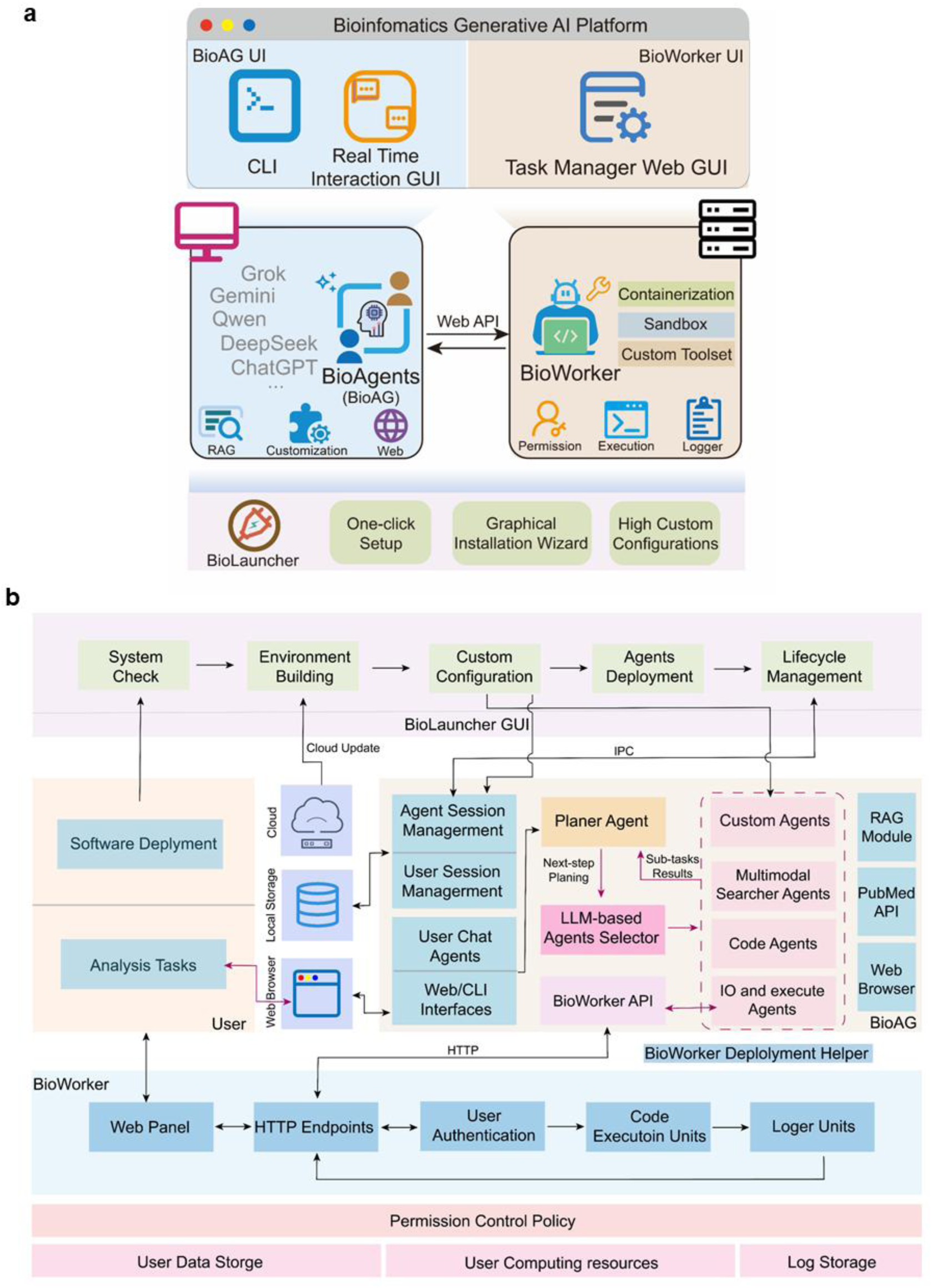
Schematic overview of the BioGAIP framework. (a) Main architecture of the BioGAIP, The BioGAIP comprises several decoupled components: BioLauncher, BioAG and BioWorker. BioLauncher is a wizard that enables graphical deployment of BioGAIP (bottom row), BioAG can adapt to a wide range of LLM API providers and support optimized agents for various bioinformatics tasks (middle left panel). BioWorker is an execution proxy with permission control, logger, and customizable functionalities, which communicates with BioAG via web API (middle right panel, see also Supplement Fig. 2). Furthermore, each component has intuitive, web-based graphical user interface for convenient usage (top row). (b) Operational details and workflow of the BioGAIP. BioGAIP provides a comprehensive, end-to-end solution. BioLauncher offers a one-stop deployment wizard and performs complete lifecycle management for BioAG. BioAG features a built-in, customizable agent system and cooperates with BioWorker to execute analytical tasks. Importantly, designed with a security-oriented architecture, BioWorker implements robust task management, logging mechanisms and permission controls to protect users’ computational assets. BioWorker can be deployed via the BioWorker Deployment Helper, a module integrated into the graphical user interface.

**Fig. 2:**
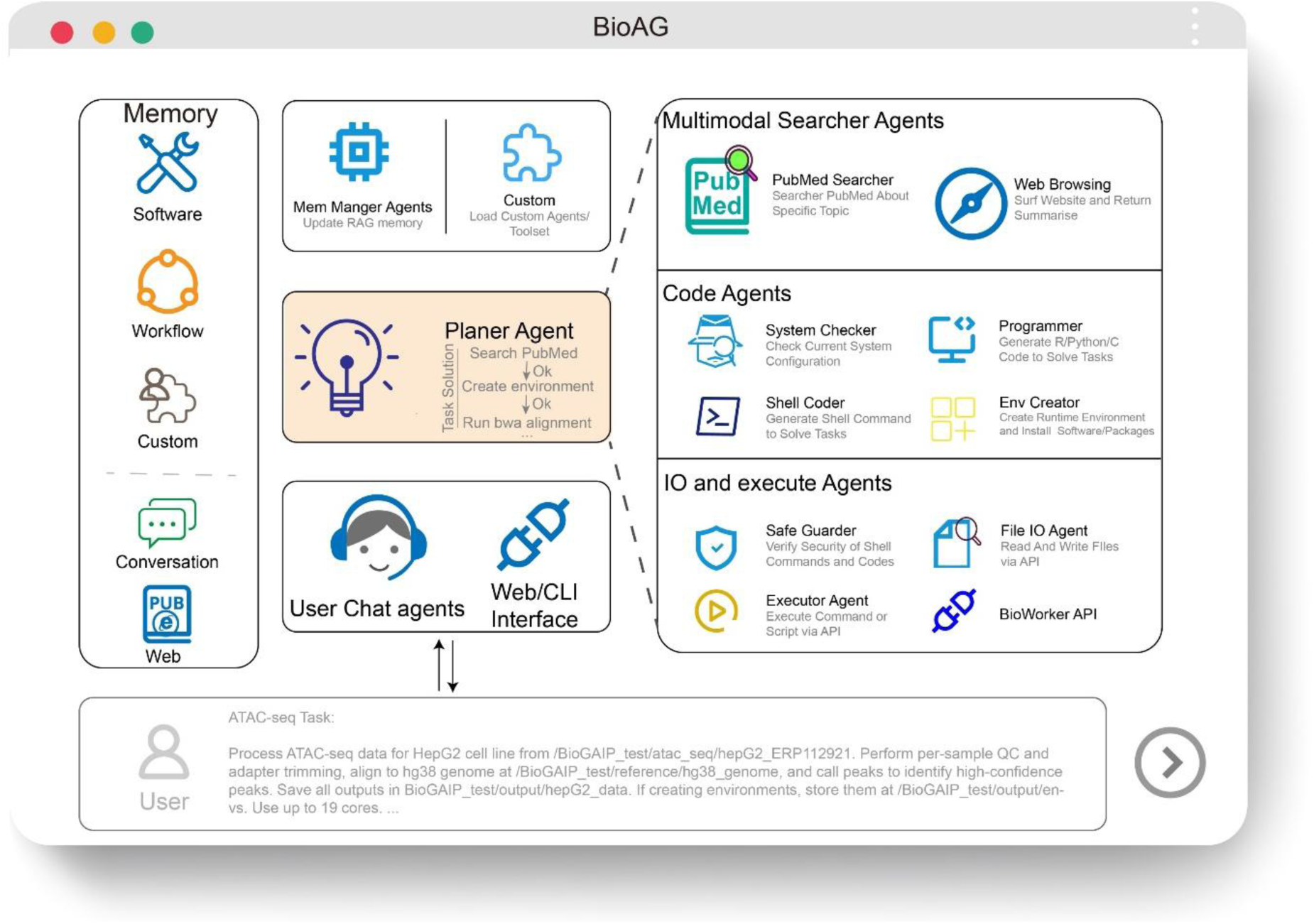
Overview of BioAG. BioAG is an optimized build-in multi-agent system. By default, the workflow begins with a user-provided task prompt submitted via web UI or command-line interface (CLI), which is then successively processed by successively processed by series of agents (such as user chat agents and planner agent). Agents leverage the user’s prompt along with external information sources such as RAG and web. The planner agent progressively plans sub-tasks, which are assigned for completion by specific agents. The remaining agents are categorized into three main groups: (1) Multimodal Searcher Agents, designed to retrieve information from PubMed and web sources; (2) Code Agents, comprising roles tailored for diverse programming capabilities (e.g., a System Checker for rapid system environment assessment, and a Programmer for writing Python or R scripts); and (3) I/O and Execution Agents, which cooperate with BioWorker to handle file operations and execute code. I/O agents also feature a dedicated Safe Guarder agent to audit code security.

**Table 1.**
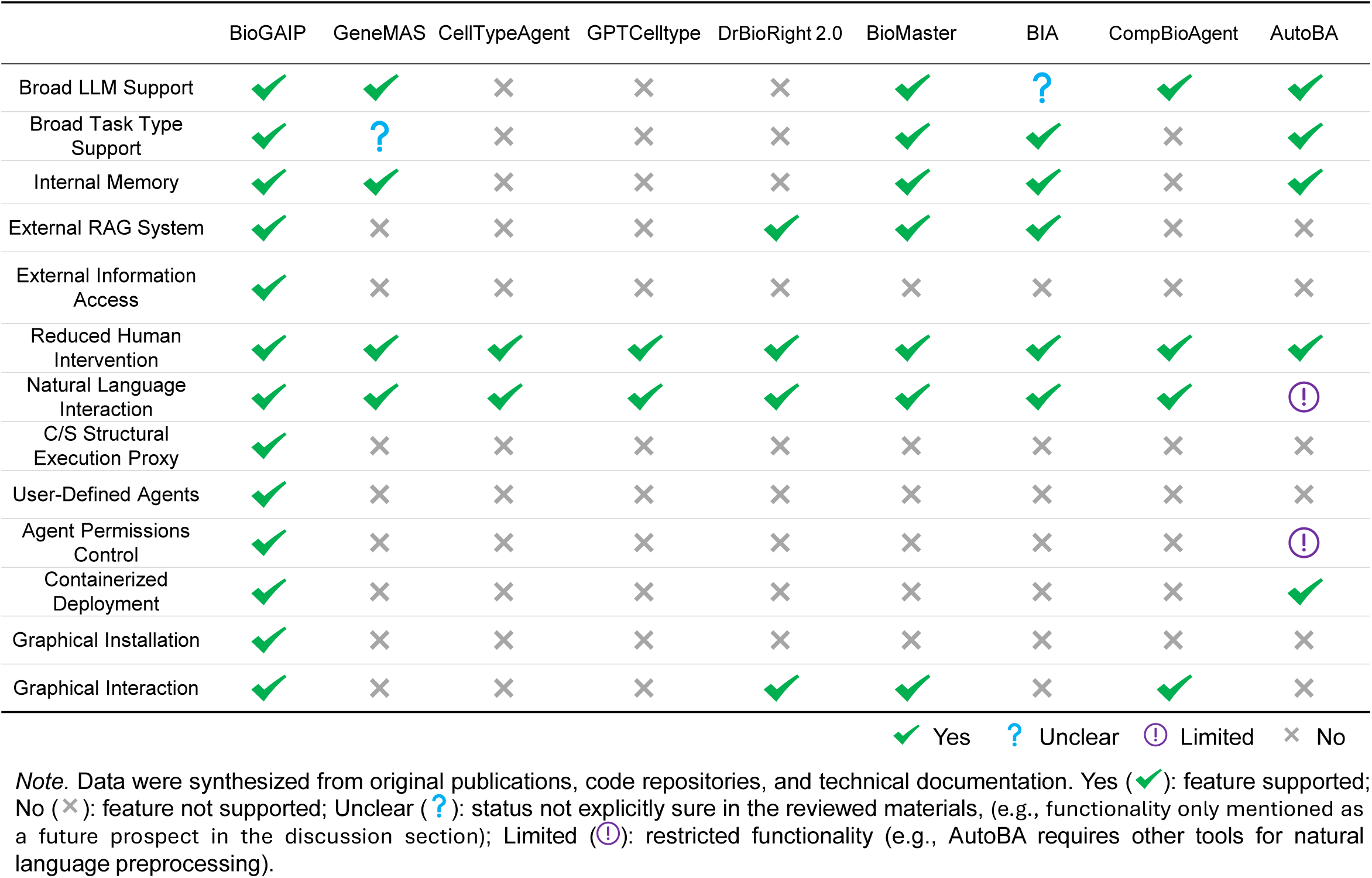
A Comparison of Bioinformatics Agent System.

|  | BioGAIP | GeneMAS | CellTypeAgent | GPTCelltype | DrBioRight 2.0 | BioMaster | BIA | CompBioAgent | AutoBA |
| --- | --- | --- | --- | --- | --- | --- | --- | --- | --- |
| Broad LLM Support | ✔ | ✔ | ✖ | ✖ | ✖ | ✔ | ? | ✔ | ✔ |
| Broad Task Type Support | ✔ | ? | ✖ | ✖ | ✖ | ✔ | ✔ | ✖ | ✔ |
| Internal Memory | ✔ | ✔ | ✖ | ✖ | ✖ | ✔ | ✔ | ✖ | ✔ |
| External RAG System | ✔ | ✖ | ✖ | ✖ | ✔ | ✔ | ✔ | ✖ | ✖ |
| External Information Access | ✔ | ✖ | ✖ | ✖ | ✖ | ✖ | ✖ | ✖ | ✖ |
| Reduced Human Intervention | ✔ | ✔ | ✔ | ✔ | ✔ | ✔ | ✔ | ✔ | ✔ |
| Natural Language Interaction | ✔ | ✔ | ✔ | ✔ | ✔ | ✔ | ✔ | ✔ | ⚠ |
| C/S Structural Execution Proxy | ✔ | ✖ | ✖ | ✖ | ✖ | ✖ | ✖ | ✖ | ✖ |
| User-Defined Agents | ✔ | ✖ | ✖ | ✖ | ✖ | ✖ | ✖ | ✖ | ✖ |
| Agent Permissions Control | ✔ | ✖ | ✖ | ✖ | ✖ | ✖ | ✖ | ✖ | ⚠ |
| Containerized Deployment | ✔ | ✖ | ✖ | ✖ | ✖ | ✖ | ✖ | ✖ | ✔ |
| Graphical Installation | ✔ | ✖ | ✖ | ✖ | ✖ | ✖ | ✖ | ✖ | ✖ |
| Graphical Interaction | ✔ | ✖ | ✖ | ✖ | ✔ | ✔ | ✖ | ✔ | ✖ |

✔ Yes ? Unclear ⚠ Limited ✖ No
*Note.* Data were synthesized from original publications, code repositories, and technical documentation. Yes (✔): feature supported; No (✖): feature not supported; Unclear (?): status not explicitly sure in the reviewed materials, (e.g., functionality only mentioned as a future prospect in the discussion section); Limited (⚠): restricted functionality (e.g., AutoBA requires other tools for natural language preprocessing).

## Results

### 1. Overview of BioGAIP

While large language models (LLMs) demonstrate broad capabilities across diverse tasks, and exhibit remarkable potential in addressing complex task, conventional prompting strategies often fall short in complex reasoning, domain-specific expertise, and specialized knowledge essential for advancing scientific discovery^8,19,20^. This limitation highlights the value of multi-agent AI systems, in which distinct LLM-based agents, each endowed with specialized roles and attributes, engage in collaborative interactions to simulate advanced reasoning processes and collectively pursue shared goals^21^. Capitalizing on this paradigm, we built BioGAIP, a comprehensive AI agent platform that enables intuitive, highly customizable, natural language-driven bioinformatics data analysis by integrating specialized knowledge and analytical functions. BioGAIP comprises a customizable built-in agent system (BioAG), an execution proxy (BioWorker), an out-of-the-box deployment assistant (BioLauncher) and their user interface (Fig. 1a-b).

BioAG incorporates a flexible agent system optimized for a broad spectrum of bioinformatics tasks through prompt engineering and in-context learning (ICL)^22^, while allowing extensive user customization of both agents and tools. The agent system supports the integration of additional knowledge bases and toolsets to enhance its capabilities and minimize errors. BioAG can also dynamically retrieve domain-specific expertise from PubMed or the internet based on the LLM assessment of the ongoing task. The framework is compatible with a wide range of LLM providers, including but not limited to ChatGPT, Gemini, DeepSeek, Qwen and Grok, and enables cost-effective role-specific model assignment (Supplemental Table 1). A key distinction from platforms like SciToolAgent is BioGAIP’s foundational skill-oriented design, each component of BioAG is optimized for the essential skills required by bioinformaticians (e.g., code writing and analysis environment construction) rather than for pre-defining task-specific tools (Fig. 2). This skill-centric paradigm renders BioAG inherently generalizable to a broad spectrum of tasks, including those not explicitly anticipated during its design, without requiring additional customization or adjustment. However, through built-in custom interfaces (detailed in Methods), it also enables the utilization of user-defined toolsets if needed, although its autonomous dependency resolution renders such custom toolsets largely unnecessary in most cases.

Given that bioinformatics analyses typically involve complex, interconnected steps, BioGAIP consolidates all agent sessions into a shared context window to facilitate optimal decision for each agent role. As observed in prior studies, LLM hallucinations accumulate with increasing conversation length, potentially disrupting task progression^23–25^. Hallucinations have also been shown to generate unsafe code, which may damage user data and assets^16, 41^, To address this risk, BioGAIP introduces a multi-layered solution. At the execution level, code evaluation is fully delegated to BioWorker, an independent, security-hardened execution proxy with user-configurable permission controls via containerization or sandbox. Architecturally, BioAG and BioWorker are fully decoupled and operate on a client-server (C/S) model through web application programming interface (API) (Supplemental Fig. 1), ensuring that BioGAIP possesses no direct code execution capabilities outside of BioWorker. At the agent level, a dedicated Security Guard Agent performs pre-execution safety checks, rejecting risky code based on the agent’s knowledge and predefined rules. Finally, to minimize hallucinations, BioGAIP integrates Retrieval-Augmented Generation (RAG), role-dependent context constraints, and content summarization strategies. These mechanisms are comprehensively detailed in the Methods section.

### 2. BioGAIP Serves as a Low-Barrier and User-Friendly Bioinfomatics Agents Platform

Designed for maximal accessibility, BioGAIP is largely cross-platform compatible, with its core components (BioAG and BioLauncher) operable across Windows, Linux, and macOS. For Windows users in particular, BioLauncher offers pre-packaged standalone executables that enable full graphical system configuration without any expertise knowledge. Compatibility testing results for BioGAIP are available in Supplemental Table 1.

Although BioGAIP supports command-line configuration and interaction, we introduced comprehensive graphical wizards and interfaces for each of its major modules to substantially lower the access barrier and enhance operational usability (Fig. 3a). These guided interfaces simplify the entire setup process and include the BioLauncher GUI, BioAG GUI and BioWorker graphical panel.

**Fig. 3:**
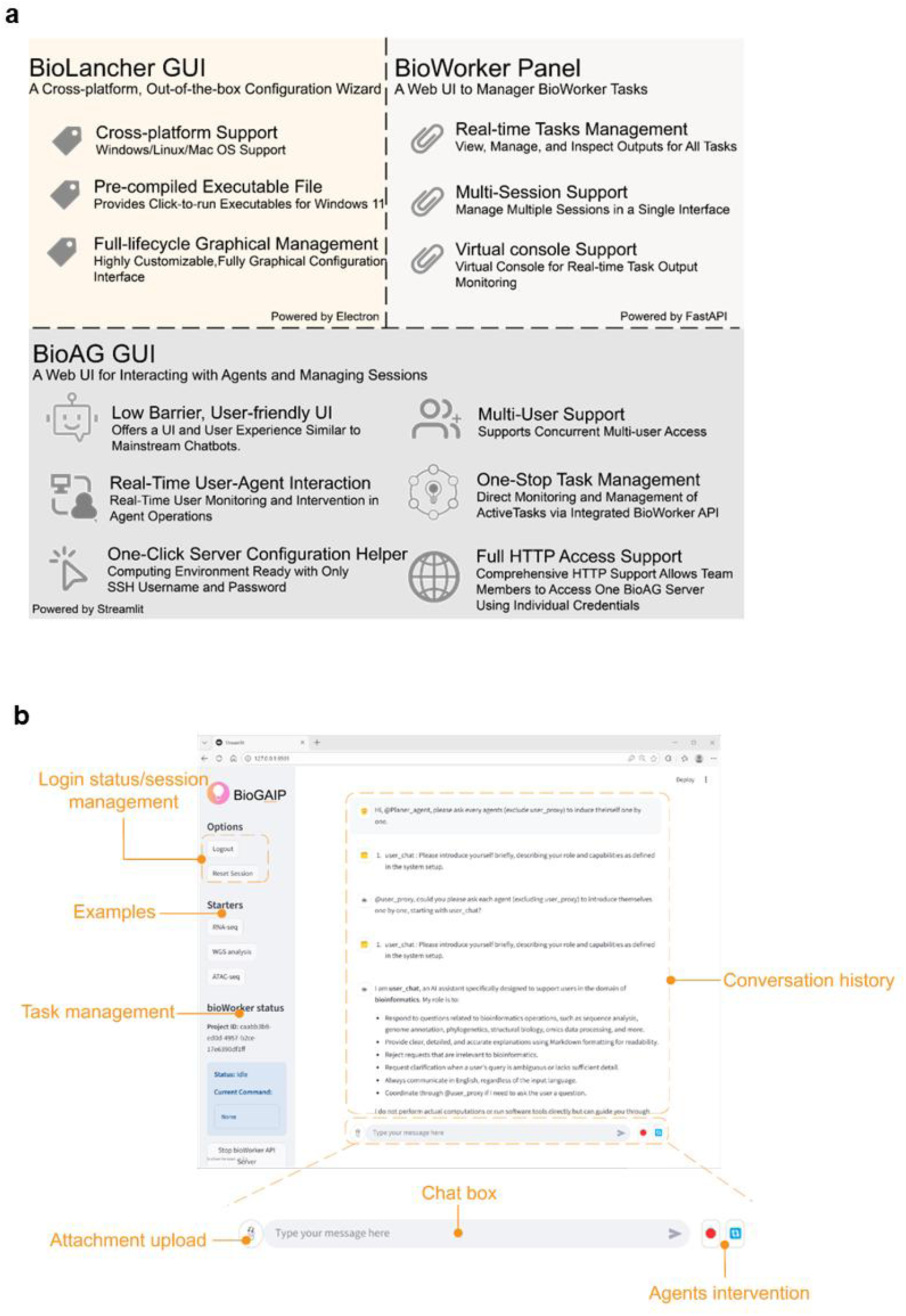
BioGAIP has a comprehensive graphical user interface for end-to-end workflows. (a) Introduction to the main features of BioGAIP’s three graphical interfaces, including BioLauncher GUI, BioAG GUI and BioWorker graphical tasks management panel. (b) Screenshot of the BioAG GUI, which provides real-time user-agents interaction and task management. The BioAG GUI consists of a collapsible sidebar and a main session area. The sidebar has quick-access management tools and task prompts examples. The session area features an input chat box, displays the real-time interaction history between the user and agents, and allows users to upload attachments and manage the execution status of the working agents.

BioLauncher is an out-of-the-box configuration wizard for BioGAIP. it can be packaged as a standalone executable file using Electron framework, bundling the web interface, essential dependencies, and scripts. This enables seamless deployment on supported operating systems (e.g., Windows 10/11), where users can initiate the wizard via a single executable file, simplifying software installation processes, thereby reducing user onboarding barriers.

BioLauncher features three principal interfaces corresponding to key configuration steps: **1. System Check**, this module verifies whether the user’s environment meets the requirements. If discrepancies are identified, BioLauncher provides targeted troubleshooting recommendations. **2. System Configuration,** Users configure their preferred LLM API providers, with broad support for commercial options, such as Qwen, DeepSeek, Gemini, Grok and ChatGPT, as well as local LLM models. Users can also integrate multiple LLMs simultaneously for hybrid workflows, enabling flexible combination based on task requirements. Advanced users may customize the agent team at this stage. Additionally, users can input external knowledge bases (e.g., custom local pipelines and reference literature) and apply custom settings. **3. Parameter Confirmation,** In the final step, users review and confirm configurations before launching BioAG. Upon successful initialization, BioLauncher seamlessly transitions the user to the BioAG interface for any remaining setup.

By default, the BioAG UI is constructed using Streamlit framework. Its operation relies on communication with BioWorker; Upon launch, users are prompted to provide BioWorker connection parameters, including the API URL and secret key. If BioWorker is not yet configured, BioAG can automate this process by requesting the user’s server IP address, username, password, and user-specified access control policies. All sensitive credentials are processed locally and not transmitted to the third parties.

Once connected to BioWorker, BioAG directs users to the BioAG conversation UI, which consists of two parts (Fig 3b). The left sidebar includes a control panel for managing session states and offering example tasks, as well as a mini, embedded BioWorker monitoring interface for overseeing the BioWorker’s operational status in real time and quickly managing tasks. The central conversation workspace displays the complete interaction history, encompassing user prompts, agent responses, and command execution outputs. Positioned at the bottom of this workspace is a chat box for submitting new prompts, accompanied by shortcut buttons for quick agent-state control. (Fig 3b).

As a security-first execution proxy, BioWorker features robust permission control, web API and user authentication, BioWorker primarily interacts with BioAG through API and operates independently (Supplemental Fig. 1, detail in Methods). However, it also provides a web-based panel for users to inspect tasks and statuses, as well as intervene manually. The panel includes key modules such as authentication, project management, web-based command interface, task monitoring dashboard, and virtual console.

Via the BioWorker panel, users can review historical task execution logs and monitor real-time task progress through a web browser, the virtual console streams live output for each command. The interface also enables manual task management and submission of new tasks. Manual interventions could automatically be detected by the agents, ensuring synchronized notification across the system.

### 3. Applicability of BioGAIP to Diverse Bioinformatics Analysis Tasks

To assess the capability of the BioGAIP system to address real-world scientific challenges, we constructed a diverse array of tasks using published datasets, including those derived from bulk RNA sequencing (RNA-seq), Assay for transposase-accessible chromatin-seq (ATAC-seq), chromatin immunoprecipitation followed by sequencing (ChIP-seq), whole genome sequencing (WGS) and single-cell RNA sequencing (scRNA-seq). Beyond these canonical paradigms, we recognized the potential interest of users in niche subdomains; accordingly, we evaluated BioGAIP’s extensibility through illustrative cases, including intronic polyadenylation (IPA) analysis via IPAFinder^26,27^, alternative transcription start site (TSS) profiling using DATTSS^28^, and cell senescence evaluation by human universal senescence index (hUSI)^29^.

During the evaluation process, we reviewed BioGAIP’s procedure and output against expert human and established publication guidelines, and verified whether tasks correct progressed to predefined endpoints. For example, generating a differentially expressed gene (DEGs) list in RNA-seq workflows. Detailed task specifications and tailored prompts are elaborated in Supplementary Material 1. Given BioGAIP’s compatibility with diverse LLM providers, our assessments spanned prominent API, including Qwen, Grok, Gemini, and DeepSeek. The foundational ethos of BioGAIP emphasizes fully automated workflow orchestration with minimal human intervention. We thus classified empirical results into three discrete tiers: (1) One-shot Success, the task was completed successfully at the first attempt; (2) Few-shot Success, success at retry. The task required retries but was completed successfully within ≤3 independent attempts. (3) Fail, No acceptable results were obtained in any of the three attempts, even with active human Intervention.

Strikingly, BioGAIP demonstrated robust performance across the assayed tasks, achieving one-shot success in most cases and autonomously overcoming analytical challenges with minimal user input (Table 2). BioGAIP does not assume any pre-installed analysis software, it automatically handles the tedious process of analysis environment installation and troubleshooting in our evaluations. For instance, during the Grok-powered RNA-seq and DATTSS analysis, BioGAIP automatically identified fault point and switched to alternative solutions when genome fasta file was unavailable. Similarly, when faced with disruptions in DESeq2 analysis, it initiated automated troubleshooting and successfully recovered (Fig. 4a-b).

**Fig 4.**
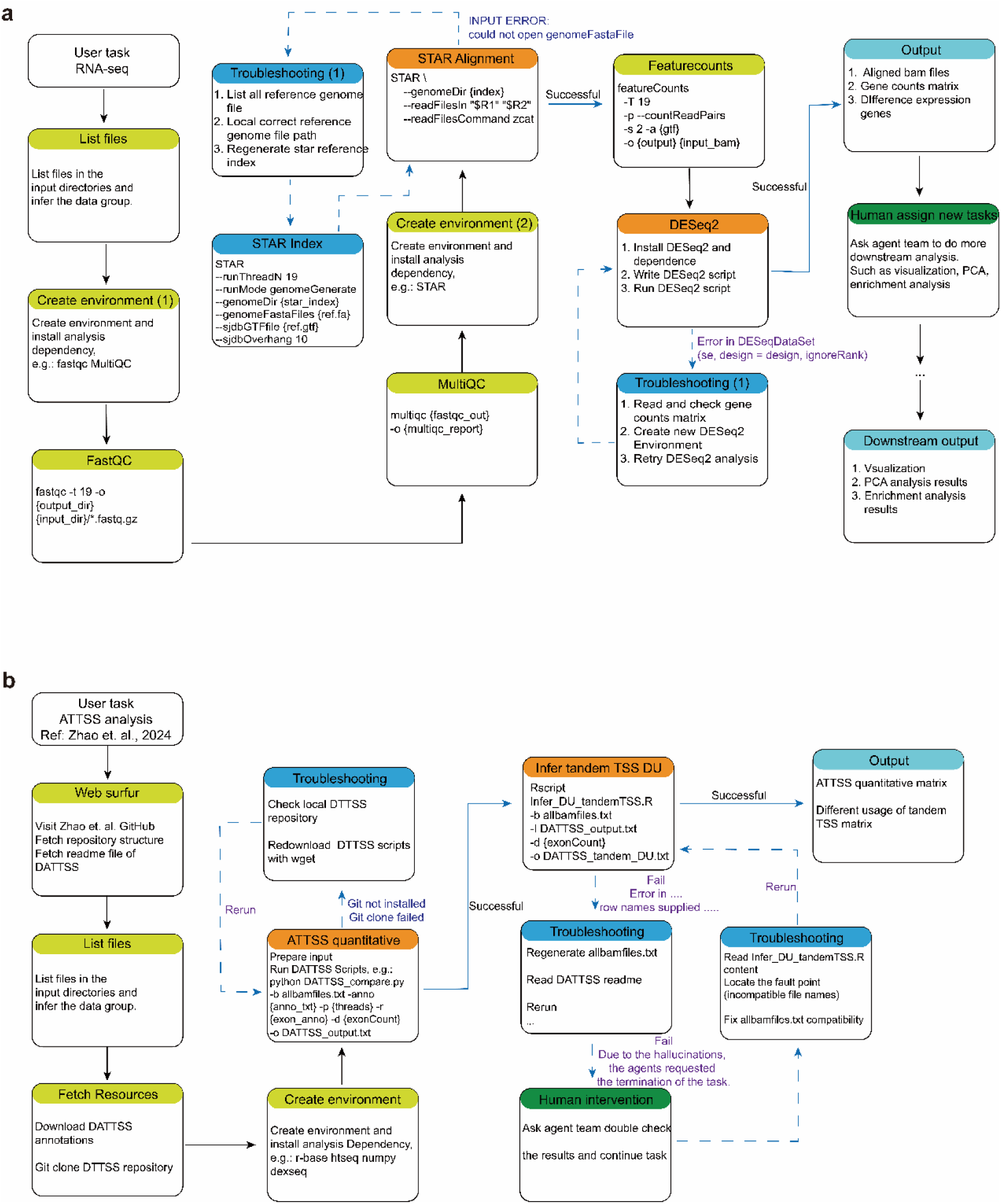
Workflow Examples illustrating applying BioGAIP to diverse omics analysis tasks. (a) Flowchart of BioGAIP analyzing RNA-seq data. The expected endpoint of this task is the generation of a list of differentially expressed genes (DEGs). During the successive process, human intervention can be introduced to prompt BioGAIP for specific downstream analyses. (b) Flowchart of BioGAIP analyzing ATTSS events. The endpoint of this task is to generate a list of differential ATTSS events. The LLM model employed in the analysis processes for (a) is qwen3-max and for (b) is grok-4-fast-reasoning. Titles in green boxes indicate normally executed steps. Titles in brown boxes indicates steps where BioGAIP encountered an error. Titles in blue boxes indicate troubleshooting steps performed by BioGAIP, with blue text showing the specific error cause(s) (when present). Titles in dark green boxes indicate steps that involved manual prompt input by the user. Arrows denote the direction of the workflow, troubleshooting flows are shown with dashed lines.

**Table 2.**
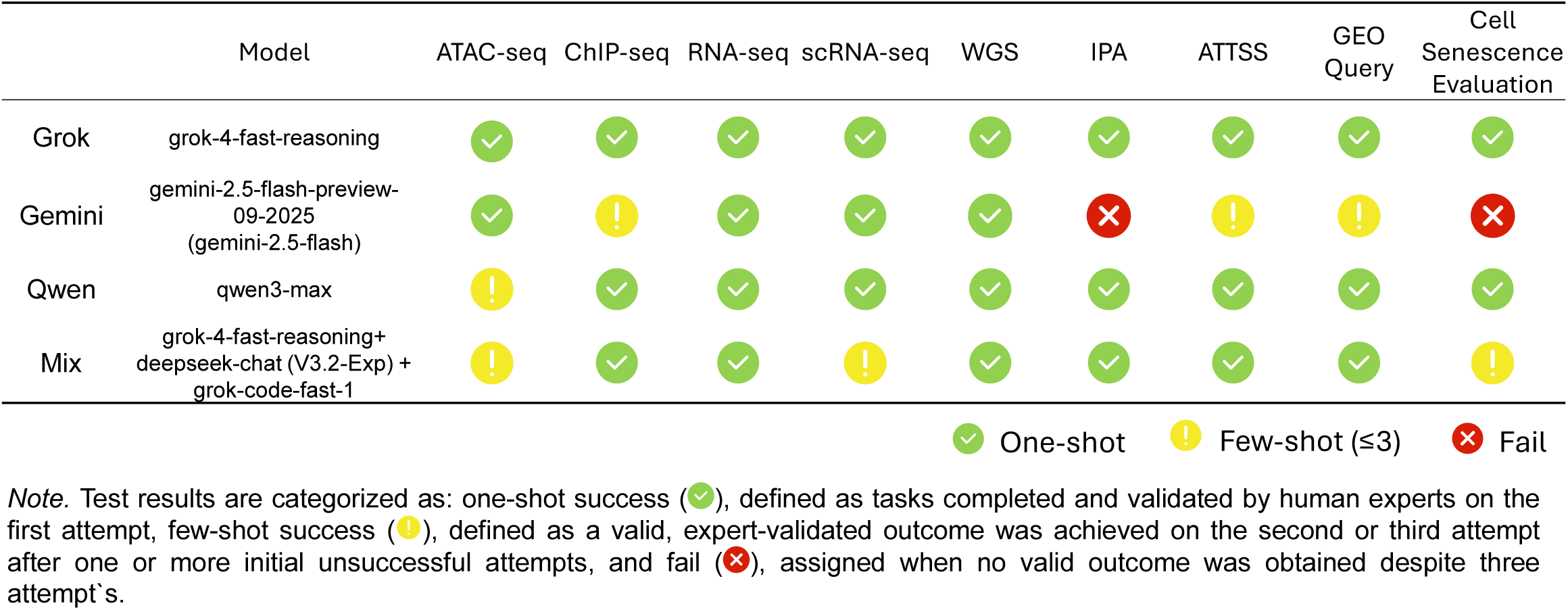
Performance comparison of different LLMs integrated within BioGAIP.

### 4. Case Study: BioGAIP Validates Key Drivers of SCLC Metastasis Through Multi-omics Analysis

To validate BioGAIP’s capacity for multi-omics analysis within real-world bioinformatics contexts, we utilized associated, published datasets encompassing bulk RNA-seq, ChIP-seq, ATAC-seq, and scRNA-seq. Our goal was to ascertain whether BioGAIP could faithfully reproduce the key biological findings of the original study. Small cell lung cancer (SCLC) is an aggressive malignancy distinguished by its remarkable metastatic potential, with most patients presenting widespread metastasis at early stages^30^. However, the molecular mechanisms underlying this exceptional metastatic capacity remain poorly defined. Previous work by Kawasaki et al. identified the *ASCL1*–*FOXA2* axis as a critical regulator of multiorgan metastasis in SCLC^31^. Here, we leveraged the same datasets originally generated by the Kawasaki et al. to evaluate whether BioGAIP could independently recapitulate these key insights using an automated agent-based multi-omics analysis.

In our evaluation, BioGAIP was provided with no access to the original literature. The agent-based system automatic determined analytical steps and parameters. Differential expression analysis correctly pinpointed significant *FOXA2* upregulation in metastasis-associated samples versus non-metastatic controls (Supplementary Fig. 2a). This finding was corroborated by single-cell RNA-seq analysis, which independently mapped high *FOXA2* expression specifically to epithelial subpopulations within tumor tissues (Supplementary Fig. 2b, Supplemental Fig. 3a). Bulk RNA-seq analysis also independently identified key genes co-expressed with *FOXA2* within the gene sets defined by Kawasaki et al., including *PROX1*, *ID2*, and *ASCL1* (Supplementary Fig. 2c). Furthermore, BioGAIP extended this insight through ChIP-seq analysis, revealing marked ASCL1 enrichment at the regulatory regions of both *FOXA2* and *PROX1* in H1836 and SHP77 cell lines (Supplementary Fig. 2d-e). Additionally, chromatin accessibility at the *FOXA2* locus was evident in all three *ASCL1*^+^ *FOXA2*^+^ Patient-Derived Xenograft (PDX) tumors, yet absent in the *ASCL1*^+^ *FOXA2^-^* PDX tumor, aligning with findings from Kawasaki et al (Supplementary Fig. 2f). The concordance across these pivotal findings underscores BioGAIP’s proficiency in diverse multi-omics analyses with high fidelity while reliably uncovering analogous biological insights.

### 5. Case Study: Discovery of Alternative Transcription Initiation Events Associated with SCLC Metastasis Using BioGAIP

To demonstrate BioGAIP’s potential in enabling novel scientific explorations, we also conducted a reanalysis on SCLC RNA-seq data from Kawasaki et al. Alternative transcription initiation of transcription start sites (ATTSS) represents an important regulatory mechanism in gene expression^32–34^. Prior studies have shown that ATTSS contributes to tumorigenesis, yet its role in tumor progression^35–37^, particularly whether aberrant ATTSS correlates with SCLC metastasis, remains unclear. We supplied BioGAIP with references to other published ATTSS analysis pipeline (publication title and DOI)^28^ and explicitly designated DATTSS as the analysis software. In most cases, BioGAIP autonomously retrieved the relevant information from the internet, installed corresponding software from GitHub, and independently set up the workflow steps and parameters to execute the full analysis. We analyzed the final outputs of BioGAIP when employing grok-4-fast-reasoning models (Fig. 4b). Interestingly, this analysis identified 200 significantly altered ATTSS events, of which 3 (1.5%) resulted in 5′ UTR shortening and 197 (98.5%) in 5′ UTR lengthening (Fig. 5a, Supplemental Fig 4a). This indicates that global 5′ UTR lengthening occurred in the primary lesions of non-small cell lung cancer with metastasis. Among genes with differential ATTSS events, only 6.5% also significantly differential in gene expression levels (Fig. 5b). Considering that changes in ATTSS can alter the coding sequence of the downstream open reading frames (ORFs) and in turn leading to biological or pathological effect. Intriguingly, we manually examined this aspect and found that only 41.2% of the ATTSS changes affected the gene’s coding sequence, suggesting that these ATTSS changes may exert effects through additional layer (Fig. 5c).

**Fig 5.**
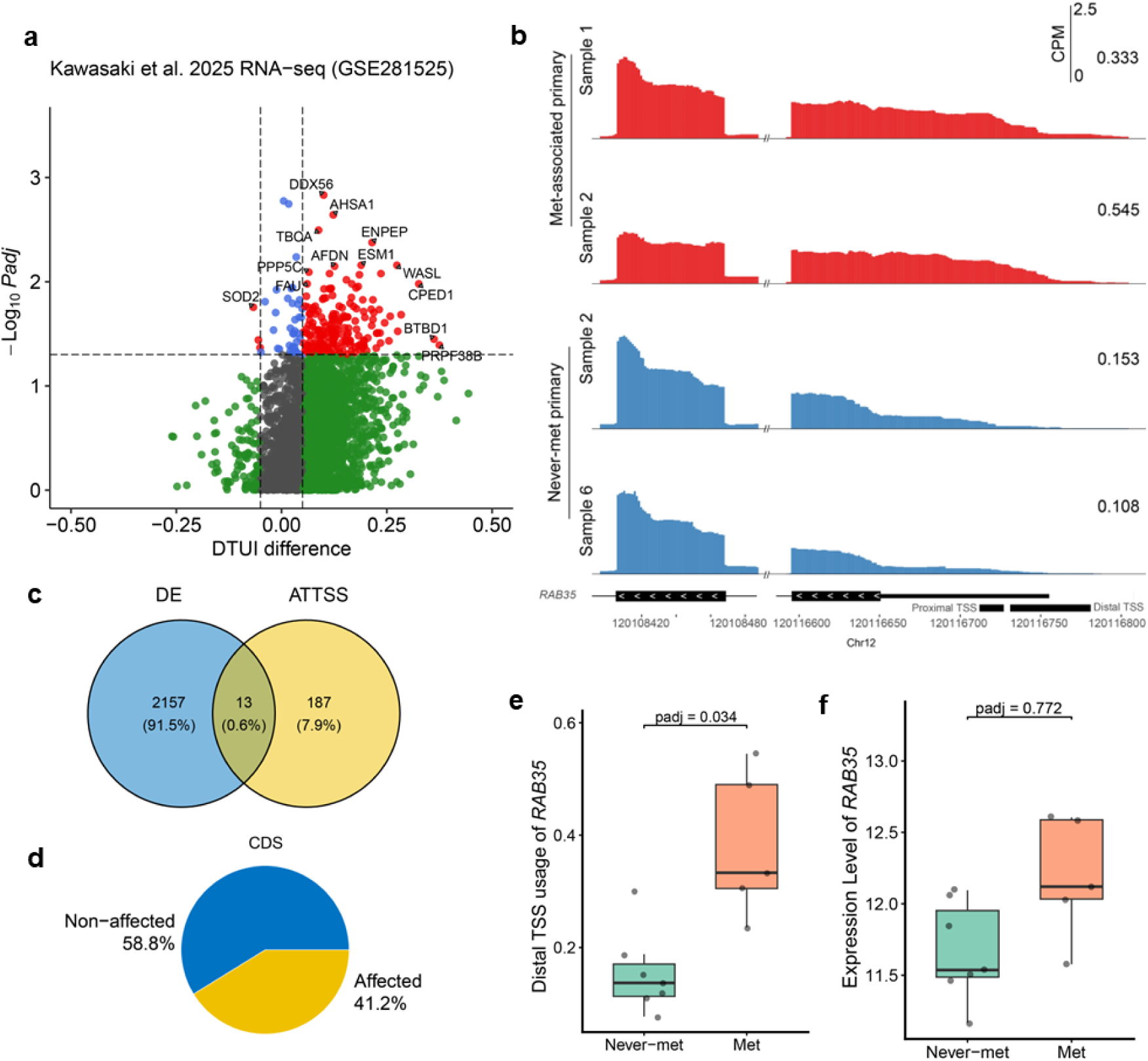
The extensibility of BioGAIP for novel bioinformatic analysis, as exemplified by Identifying Different Transcription Initiation Events Associated with SCLC Metastasis. (a) Volcano plot of differential ATTSS events. (b) RNA-seq density plots showing the high distal TSS usage of *RAB35* in met-associated primary tumors and never-met primary tumors. (c) The Venn diagram between different ATTSS events and differential expression genes. (d) The effect of different ATTSS events on gene coding region. (e) The comparison of distal TSS usage of *RAB35* between never-met primary and met-associated primary tumor. (f) The comparison of gene expression of *RAB35* between never-met primary and met-associated primary tumor. Statistical significance of distal TSS usage was assessed using the DATTS^28^, statistical significance of gene expression was assessed using the DESeq2^49^. The analysis of (c) was completed by human experts based on the output of BioGAIP.

For example, TSS usage of *RAB35* and *AP4E1* ranked among the top 30 by significance of differential tandem TSS usage in showed significant difference between metastatic and non-metastatic SCLC (Fig. 5d,e, Supplemental Fig. 4b,c). It’s 5′ UTR difference did not affect the coding region of *RAB35* (data not shown), and no significant difference in both *RAB35* and *AP4E1* expression levels was observed between metastatic and non-metastatic SCLC primary tumors (Fig. 5d, Supplemental Fig. 4d).

To further validate this finding, we applied BioGAIP to analyze an independent SCLC dataset (GSE60052), which includes 7 normal tissue samples and 79 tumor samples. Consistent with the previous results, *RAB35 and AP4E1* showed no significant expression difference between normal tissues and SCLC tumors in this dataset (Supplemental Fig. 4e,f).

## Discussion

BioGAIP automates multi-omics exploration by harnessing diverse LLM APIs and optimized agents to perform complex analysis tasks. Specifically, BioGAIP allows users to execute end-to-end workflows using simple one-shot or few-shot prompts, thereby eliminating the need for programming expertise. Several prior works have achieved notable advancements in this area, underscoring the pivotal role of LLMs in facilitating biological research^11,15,38,39^. This is particularly evident in bioinformatics, where task-oriented solutions such as DrBioRight 2.0, GPTCelltype, and CellTypeAgent, alongside general-purpose agent systems (such as AutoBA and BioMaster), have demonstrated impressive capabilities. To the best of our knowledge, BioGAIP surpasses existing approaches by incorporating additional enhancements that broaden its applicability and efficiency (Table 1).

Many similar methods rely on specific platforms (e.g., dedicated websites) or restricted execution environments, and typically offer limited LLM provider support, thereby raising the barrier to entry and restricting software accessibility. In this study, we implemented cross-platform support for BioGAIP, developed a fully graphical interface for installation and interaction, validated compatibility with most mainstream LLM APIs, and provide a plug-and-play executable for Windows, ensuring accessibility to the broadest audience. notable, BioGAIP is designed to permit extensive customization, allowing advanced users to adapt BioGAIP for specialized and personalized research applications. To enhance security and reliability, we incorporated BioWorker, a secure and isolated execution proxy that leverages client-server architecture and containerization for flexible deployment across diverse computational environments.

In contrast to agentic frameworks dependent on static global planning, BioGAIP implements a default reactive approach, enabling agents to collaboratively identify and execute actions in a stepwise manner. This reactive strategy ensures that each planning step is dynamically conditioned on intermediate results accumulated from preceding actions. We evaluated BioGAIP across a diverse collection of public datasets and challenging real-world tasks, and empirically observed that it successfully completes these complex tasks in a one or few shot prompts. These results highlight the system’s strong performance in problem comprehension and self-correction.

While the results presented primarily highlight BioGAIP’s utility within limited, proof-of-concept scopes, its applicability extends significantly further. In our ongoing internal testing and user feedback, BioGAIP has already demonstrated high performance in broader contexts, including WGS, automated retrieval of large-scale GEO datasets, omics data curation, and other tasks. We will continuously expand applications examples, including new use cases, tutorials, and prompts, and regularly updated on the BioGAIP notebook website which will be made publicly available upon the publication of this article (https://notebook.biogaip.top).

Nevertheless, there still remains room for further development to expand BioGAIP’s capabilities. Despite the integration of multimodal web access, its ability in multimodal interpretation, such as image interpretation, still remains limited. Interactions with complex datasets, particularly omics data, still primarily rely on text-based input/output. Ongoing advances in multimodal LLMs and agent frameworks are expected to overcome these shortcomings. Additionally, context length accumulates rapidly in extended sessions, which can exacerbate hallucinations, increase latency, and elevate token costs. Although per-agent context limits, content-summarization mechanism, RAG and recent improvements in commercial LLM context windows length provide some relief (detail in Methods), BioGAIP does not currently include additional mitigation strategies such as selective summarization or context compression. Future iterations could introduce optional, user-configurable context-management techniques to mitigate these issues.

Another challenge lies in the reproducibility of analyses, which stems from the inherent stochasticity of LLM outputs. As with many comparable tools, even when identical prompts are used, obtaining fully consistent results across independent runs remains challenging. To mitigate this limitation, users have the option of employing the browser’s built-in save functionality to archive and share complete session conversation histories. In addition, BioWorker provides a query interface that lists every historical command executed within a project. For advanced users, BioGAIP offers an experimental feature to export the project configuration file; under the same software version, this file can be imported on other devices to restore the session state.

Despite these ongoing challenges, BioGAIP provides a robust foundation for automating complex multi-omics analysis workflows. Future efforts will focus on integrating comprehensive multimodal capabilities, more rigorous hallucination mitigation, and seamless session sharing. Ultimately, BioGAIP showcases the potential of LLM-driven agents to streamline and empower data-driven biological discovery, making sophisticated analyses accessible to a broader audience.

## Methods

### 1. BioAG is a Multi-agent System

Modern bioinformatics tasks require not only the processing of large-scale data but also the flexibility to support continuous hypothesis evolution. To meet this need, BioGAIP integrates BioAG, an agentic framework built on AutoGen framework^18^ that utilizes prompt engineering and in-context learning (ICL) with LLMs. Our selector group paradigm facilitates numerous specialized agents operating under the "Do one thing and do it well" principle. We systematically organize these agents into following six main groups (Fig. 2):

#### User Chat Agents

This is the user interaction layer of BioAG, responsible for understanding user queries and conducting dialogue. Its architecture, which combines an LLM agent with UserInputProxy function, provides a unified entry point that allows the entire system to be Integrated with multiple front-ends like command line interface (CLI), or streamlit-powered graphical user interface (GUI).

#### Multimodal Retrieval Agents

To bridge the gap between pre-training LLMs and real-time information needs, our architecture features multimodal retrieval agents, composed of two specialized components. The first is a PubMed-specialized agent for targeted real-time literature retrieval via PubMed API. The second is a general web browsing agent, designed to transcend the boundaries of traditional academic databases. By programmatically accessing dynamic web platforms such as GitHub and Google Scholar, it extends the framework’s reach to capture the rapidly evolving research landscape, thereby incorporating external data and code repositories essential for bioinformatics.

#### Code Agents

This ensemble comprises specialized programming roles. It includes a system configuration agent for inspecting computing environments, an environment specialist agent for managing dependencies via micromamba and pip, a shell scripting expert that orchestrates Linux-based tool execution, and a general programmer agent that handles diverse coding tasks beyond these specialized domains.

#### IO and Execute Agents

BioGAIP is engineered for robust file system interaction and the generation of analysis outputs consistent with conventional bioinformatics workflows. To achieve this, BioAG integrates two specialized agents: file agents and execute agents. These components are optimized through prompts that leverage large language models’ function-calling capabilities to interface with the BioWorker web API. To address potential risks associated with LLM hallucinations that might generate erroneous commands compromising user assets, we implement safeguard agent as part of BioGAIP’s security framework. These agents perform pre-execution validation by screening commands against both the model’s knowledge and predefined security rules, ensuring only verified command proceed to execution.

#### Memory Management Agents and Module

BioGAIP is designed to integrate external domain expertise and user-defined knowledge bases. However, the length of such textual data may exceed the context window constraints for LLMs. To overcome this limitation, we employed a retrieval-augmented generation (RAG) method. This RAG method integrates a vector database (ChromaDB) with embedding models to enable efficient storage, indexing, and on-demand retrieval of pertinent knowledge segments^40^. This approach separates extensive knowledge storage from the model’s constrained context window, thereby enabling agents to access comprehensive, domain-specific information while mitigating substantial computational expenses. In default, most agents are allowed access to this memory system. Memory management agents maintenance and dynamic updating of the memory across the entire workflow.

#### Planner Agent

The planner agent constitutes the central component of BioAG, where we implemented the select group model integrated with the chain-of-thought (CoT) principle to accomplish task-driven data analysis. Specifically, unlike the widely adopted round robin group model, the specialized agents within BioAG are not constrained by a predefined agents execution sequence, although some specific roles, such as safety check agents, which are always executed prior to code execution. Instead, we introduced a planner agent that determines what to do next step based on the existing results and assigns appropriate roles to advance the workflow. Select group model equipped with a model-based next-speaker selection mechanism, enabling the team to analyze the current conversational context, including conversation history and participant attributes (e.g., agents name and description), to dynamically determine the most suitable agent to proceed.

Assigned as an important orchestrator, the planner agent accesses the full conversational context and system memory by defaults. It continuously evaluates task progress, assigns subsequent steps, and assesses the outcomes of executed actions. Diverging from systems such as BIA, which typically formulate an explicit, comprehensive plan at the start of a task, BioGAIP requires the planner agent to allocate and schedule only one or few tasks at a time. This design avoids early over-commitment to a rigid plan and enables dynamic evaluation and adjustment at any stage of the process. Through this step-by-step, adaptive approach, embodying the CoT paradigm and in situ learning, the system achieved enhanced complex reasoning capabilities via structured intermediate reasoning steps.

### 2. Agents Management Module for BioAG

While BioGAIP ships with a pre-optimized agent team configuration, advanced users may prefer to define custom agent configuration and LLM providers. To this end, BioAG introduces an agent management module that supports complete customization of the agent team through YAML configuration files or graphical editor. User-provided configurations are validated and transformed into AutoGen-compatible agent instances.

BioAG offers a unified interface for users to designate and configure preferred LLM API endpoints. The interface supports a broad spectrum of mainstream LLM APIs, as well as locally hosted LLMs, for instance, Ollama. Furthermore, users can assign distinct LLM providers to different agent roles within one agent team, capitalizing on model-specific advantages to improve cost-effectiveness and augment systemic adaptability.

### 3. BioWorker: A Security-Oriented Multi-Session Execution Proxy

While popular LLM-powered solutions such as GitHub copilot and ChatGPT have demonstrated utility in code generation, studies indicate they can introduce a higher rate of vulnerabilities compared to human-written code^16^. Analysis has revealed a range of flaws in LLM-generated code, including trivial errors, security-related code smells, and issues cataloged under the common weakness enumeration (CWE)^16,41,42^. The automated execution of such unverified scripts without human review and approval poses a critical threat to modern bioinformatics infrastructures, which necessitate strict security protocols due to their interaction with sensitive data and integration with physical server, costly high-performance computing (HPC) cluster and cloud-based computational resources.

To address these challenges, we designed BioWorker, a security-centric multi-session execution proxy. BioWorker operates with BioAG in a client-server (C/S) paradigm, where BioAG functions as the client and BioWorker serves as the backend server. In its present version, BioWorker implemented permission control via container-based isolation or experimental sandboxing mechanisms. It also implemented a session management that maintains isolated execution environments with specific context for each active session (Supplemental Fig. 1). This architecture enables multiple BioAG clients to connect concurrently to a single BioWorker instance without interference, significantly enhancing system flexibility and computational resource utilization. Beyond delivering API endpoints for BioAG integration, BioWorker incorporated a dedicated web-based management interface, termed the BioWorker Panel, which facilitates real-time monitoring and administration of BioWorker activities via a web browser.

The core functionality of BioWorker lies in the secure execution of commands submitted by BioAG, this component is designed to handle BioAG inputs, manage execution processes and address any errors that may arise during execution. The BioWorker process involves the six steps as follows:

(1) **Authentication:** BioWorker’s interactions are primarily based on its web API. To prevent unauthorized access, BioWorker incorporates a built-in authentication module. For any http request, BioWorker parses the *x_api_key* field from the request header section and compares this field with the API specified during BioWorker startup. For unauthorized access, BioWorker rejects the request and returns a 401 error. By default, communication between BioWorker and BioAG is plaintext. Since BioWorker’s communication is entirely based on its web API, users can simply encrypt the communication traffic using Secure Sockets Layer (SSL) to achieve higher communication security.
(2) **Project Initialization:** As an independent execution proxy, BioWorker orchestrates tasks independently for each project, upon receiving the initialization request from BioAG at session onset, it promptly launches a session manager to configure the execution environment (e.g., working directory and usable custom toolset) and instantiates task and log managers. Subsequently, BioWorker returns the project’s universally unique identifier (UUID) and keeps the session alive for its entire session lifecycle.
(3) **Input preparation:** The BioWorker uses FastAPI to fetch the required input parameters from the BioAG, including command content, command type, command lifecycle, and others.
(4) **Command execution:** Upon receipt of the command content, BioWorker automatically identifies the appropriate execution modality based on the command’s type (e.g., shell scripts, Python, or R scripts). All command executions are handled asynchronously to ensure the main thread against blocking or interruption. Once the task begins execution, BioWorker returns the task’s universally unique identifier (UUID) to BioAG via its http endpoint and output execution results after finished.
(5) **Output and Error Handling:** The executor employs inter-process communication (IPC) protocols to intercept standard output (stdout) and standard error (stderr) during command execution. Captured outputs or errors become accessible to users through the BioWorker panel. Following task completion, such logs are preserved as session history. In the current implementation, BioWorker does not send push notifications to BioAG for task status or output updates, instead, monitoring of these details is handled through BioAG’s periodic polling of http endpoints.
(6) **Task Status Query and Management Services:** The BioWorker system implements task status querying through a set of http endpoints. For any unique task UUID, upon successful authentication, BioWorker returns to the requester the task’s status (indicating whether it has completed), real-time output, execution duration, and any relevant status code. Additionally, BioWorker also supports http endpoints for managing the BioWorker and associated tasks life cycle.

### 4. External Knowledge Bases and RAG Memory

While LLMs exhibit powerful generative capabilities, their reliance on static training corpora often limits precision and induces hallucinations^43,44^. To address this, BioGAIP employs RAG, which dynamically interfaces with external repositories during inference. This approach significantly enhances factual fidelity and relevance by grounding outputs in up-to-date information, all without the computational overhead of model retraining^45^. In BioGAIP, we incorporate domain-specific assets, such as software repositories, workflow databases, and user-defined datasets, complemented by agent-retrieved public information, to form a modular RAG framework that optimizes LLM efficacy for specialized bioinformatics inquiries.

#### Bioinformatics Workflow Databases

In BioGAIP, we harness the nf-core project, a community-led repository prioritizing standardization, portability, and reproducibility in analytical workflows. Through its API and GitHub integration, we retrieve 137 high-caliber bioinformatics pipelines encompassing varied domains and data modalities. These pipelines are code in Nextflow, a workflow orchestration language. To render them compatible with LLM integration, we apply the gitingest to convert associated Nextflow repositories into streamlined, LLM-optimized textual formats. Subsequently, embeddings are computed using the all-MiniLM-L6-v2 model (or user-specificed) and stored persistently in ChromaDB to support downstream RAG applications.

#### Bioinformatics Software Databases

Given the disparate installation protocols inherent to bioinformatics software, maintaining consistent computational environments remains a significant challenge. Systems such as Conda alleviate these operational bottlenecks by utilizing recipes-based configuration files to manage dependencies and ensure reproducibility. BioGAIP integrates a parser that harvests this rich metadata from open-source conda repositories like Bioconda, subsequently encoding it into a vectorized format (e.g., ChromaDB) to facilitate retrieval-augmented generation.

#### PubMed and Web Based RAG Construction

To equip the model with real-world data robust awareness capabilities, we developed a PubMed retrieval agent and a web browsing agent (see above). When network access is available, the PubMed retrieval agent interacts with the PubMed API via an internally interface, autonomously determining query-specific keywords to retrieve relevant literature. For articles indexed by PubMed and openly accessible, the agent fetches and ingests full-text content into the ChromaDB vector database; otherwise, it accesses only publicly available metadata, such as titles and abstracts. The web browsing agent, a pre-integrated multimodal component in BioGAIP, is implemented using AutoGen’s MultimodalWebSurfer object, which launches a Chromium browser and leverages playwright for interactive web navigation and autonomous behaviors on webpage content, thereby enabling agents to perform context-aware internet exploration adapted to user queries or task requirements. Agents can decide whether to incorporate the content found through web surfer into their memory.

#### User-Customized RAG Content

BioGAIP enables users to integrate customized, domain-specific knowledge into the RAG system, encompassing curated documents, internal protocols, or other proprietary resources. Supported input formats include plain text files (e.g., TXT, MD, HTML) as well as portable document types like PDFs. For files with rich text, such as PDFs, BioGAIP extracts the textual content using third party libraries like PyPDF prior to indexing in the vector database. This on-device processing computes the embeddings locally on the user’s local machine. As a result, it limits the exposure of private information to external LLM API providers, unlike approaches that incorporate complete data into prompt contexts. This method thereby enhances the security and confidentiality of private materials and technical datasets.

### 5. Web-Based User Interface Implementation

The BioGAIP platform features an intuitive, fully graphical interface composed of three core components: the graphical software configuration wizard (BioLauncher), the BioAG user interface (UI), and the BioWorker panel (Fig. 3a,b). These interfaces are developed using HTML, JavaScript, and Python. Specifically, the BioAG UI is built with web frameworks such as streamlit to support real-time web-based interaction with LLM agents. User authentication and session management are handled by persisting session data in local JSON files. Communication between the BioAG UI and the AutoGen-driven agent team service is conducted asynchronously via inter-process communication (IPC), with the UI updating and managing the state of agent services. For BioWorker, the third-party Python library FastAPI is employed to construct its web API services and administrative panel (Supplemental Fig. 1). As a setup wizard, BioLauncher enables installation and configuration of both BioAG and BioWorker through a graphical interface, requiring no technical expertise. BioLauncher built with electron, node.js and streamlit for cross-platform compatibility. BioLauncher is also distributed as a standalone executable for Microsoft Windows 10/11.

### 6. Containerization and Sandbox Support

To ensure optimal cross-platform compatibility, distributability, security, and simplified installation, BioGAIP is constructed using containerization technologies. These technologies also control resource access for BioWorker to enhance security. The system primarily supports Docker, with experimental integration for alternatives such as Singularity. Dedicated build scripts (Dockerfiles) are provided for BioAG and BioWorker, incorporating micromamba docker images (mambaorg/micromamba) to enable robust micromamba support within BioWorker. Final images are generated locally via the docker build command, followed by conversion to Singularity Image Format (.sif) files using the singularity build command. Automated container build and deployment scripts are also distributed as part of the BioLauncher graphical interface.

In environments lacking containerization support, BioGAIP experimental permits container-free execution. In this situation, to maintain BioWorker access controls, sandbox technologies, such as Bubblewrap or Landlock (kernel-dependent), are employed for file access restriction. Sandbox activation requires launching BioGAIP from graphical interface and configuring BioWorker sandbox through the settings panel.

### 7. Fault Recovery and Hallucination Mitigation

In BioAG, we implement a session management system that organizes agent sessions, user profiles, conversation histories, and associated resources. The running states of agents are persistently stored in JSON files. In the event of a failure, if the user elects to resume the task, BioAG reconstructs the agent team and attempts to restore the previous state by invoking the *load_status* function to load data from the preserved JSON file.

As observed in prior studies, LLM hallucinations accumulate with increasing conversation length, potentially disrupting task progression^23–25^. BioAG addresses this through a multi-tiered approach. First, it integrates RAG system, a proven hallucination mitigation technique. Second, it enforces context length limits for specific roles, e.g., code agents, to preserve task focus and reduce cumulative errors. Finally, BioAG features a fallback summary-restart mechanism that can be engaged either automatically or manually. Upon triggering, the system halts the active agent team and reformats the accumulated conversation history into an LLM-friendly Markdown text, this text is then processed by an independent, lightweight agent designed specifically to review, summarize, and consolidate the contextual data. A new prompt is then generated based on the summarized results, enabling seamless task resumption with a freshen agent team.

### 8. LLM Model selection

BioGAIP supports a wide range of commercial and open-source LLM APIs. In this study, we evaluated models from multiple providers, including Qwen (qwen3-max), Gemini (gemini-2.5-flash-preview-09-2025 and gemini-2.5-flash), Grok (grok-4-fast-reasoning), and the compatibility of Qwen (qwen-plus-2025-12-01), Gemini (gemini-3-flash-preview), Grok (grok-4-0709, grok-4-fast-reasoning and grok-code-fast-1), ChatGPT (gpt-4.1-mini and gpt-4o), and DeepSeek (deepseek-reasoner) were also tested. By default, a single model is assigned to all agents in one test configuration, however, due to compatibility constraints with the Gemini API, the execution agent was configured to use deepseek-chat V3.2-Exp when testing Gemini. In addition, we also evaluated a scheme that combines the use of multiple models (aka mix model, grok-4-fast-reasoning+deepseek-chat_V3.2-Exp+grok-code-fast-1). All APIs were officially subscribed, and corresponding base URLs, API keys, and model names were configured in BioGAIP for test use.

### 9. Test Task Design

To evaluate the performance of BioGAIP, we designed analytical tasks covering diverse data types using publicly available datasets. The study utilized datasets retrieved from the Sequence Read Archive (SRA) or Gene Expression Omnibus (GEO), specifically accession numbers GSE281523, GSE281524, GSE281525, GSE281740, GSE60052^46^ and Cancer Cell Line Encyclopedia (CCLE, PRJNA523380)^47,48^ (detail in Supplemental Table 2). With the exception of GSE60052, these datasets originate from a single comprehensive study and represent a coherent multi-omics collection, encompassing RNA-seq, ChIP-seq, Assay for ATAC-seq, and scRNA-seq^31^. We selected this related series to rigorously evaluate BioGAIP’s capability to independently reproduce key findings from original publication. By assessing whether the agent could derive comparable biological insights, we aimed to validate its proficiency in solving complex, real-world biological problems.

For each task, we created similar initial prompts for agent teams powered by different LLM models, varying in input and output file paths (Supplemental materials 1). In a single attempt, the initial prompt was provided to BioGAIP to execute the analysis. Each task was allowed up to three attempts. An attempt was considered a failure if BioGAIP produced a serious error, severely deviated from the task objective, generated invalid outputs, or terminated the session without completion, excluding cases caused by unexpected network interruptions. Test results were stratified into three categories: one-shot success, defined as tasks completed and validated by human experts on the first attempt, few-shot success, defined as a valid, expert-validated outcome was achieved on the second or third attempt after one or more initial unsuccessful attempts, and fail, assigned when no valid outcome was obtained despite three attempts.

Note that early experiments utilized the gemini-2.5-flash-preview-09-2025 model to evaluate the performance of the Gemini family. This preview version was subsequently deprecated by Google and migrated to the gemini-2.5-flash model. Consequently, a subset of the Gemini-related tasks reported herein were performed using the gemini-2.5-flash model.

### 10. Data Processing and Visualization

Since most underlying LLMs currently lack native multi-modal visual processing capabilities, BioGAIP faces challenges in iteratively refining graphical outputs. While the agent can automate the generation of visualization scripts, it is unable to perceive or self-correct the resulting images. Consequently, the generated plots often fall short of publication standards unless subjected to extensive, multi-turn human prompting. To ensure aesthetic rigor and clarity, the majority of figures in this manuscript were manually generated using R (v4.4.1) and the ggplot2 package (v3.5.2).

## Data availability

The published datasets accession number included in this article are recorded in method, The GRCh38 reference genome used to align the raw sequence files was downloaded from NCBI (https://ftp.ncbi.nlm.nih.gov/genomes/all/GCA/000/001/405/GCA_000001405.15_GRCh38/seqs_for_alignment_pipelines.ucsc_ids/GCA_000001405.15_GRCh38_full_analysis_set.fna.gz). The gene annotation file was downloaded from GENCODE (https://www.gencodegenes.org/human/release_48.html). The single-cell RNA-seq reference package (refdata-gex-GRCh38-2024-A.tar.gz) from the Cell Ranger website (https://www.10xgenomics.com/support/software/cell-ranger/downloads). The other data that support the findings of this study are available from the Supplementary Information or corresponding author upon reasonable request.

## Code availability

BioGAIP be implemented in both Python and Shell, the codes are provided in a GitHub repository (https://github.com/zhangjy859/BioGAIP), which will be made publicly available upon the publication, and a Zenodo repository (doi: 10.5281/zenodo.19179297 that is currently accessible to reviewers along with a detailed usage guideline at BioGAIP notebook (https://notebook.biogaip.top). The code for constructing the RAG from external nf-core projects and non-commercial Conda channels is provided as a standalone, independent repository and can be downloaded here: https://github.com/zhangjy859/RAG_tools. We built a cloud platform to support BioGAIP deployment, but these cloud resources are independent of BioGAIP main functionalities. Owing to proprietary content, the code is available from the corresponding author upon request under appropriate agreements.

## Acknowledgements

We thank Shuwen Zhao, Limin Lin, Haijing Ma, Jingfei Zhang, Lingxian Wang, and all users who join in our internal early alpha test. We thank Dr. Peng Yu for the suggestion of artwork design. We thank Dr. Zhaozhao Zhao for the suggestions about data analysis. We especially thank Jing Wang for suggestions during the initial stages of this study. The design and launching of this study were financially supported by the National Key Research and Development Program of China (2023YFC3603300, awarded to T.N.).

## Author contributions

J.Z. and T.N. conceived of the project. J.Z. designed and wrote the initial version of the software. J.Z. and P.G. designed and performed the data analysis and evaluation pipeline for the project. J.Z. improved subsequent versions of the software with assistance from G.J. Additionally, J.Z., P.G., G.J., M.Z., G.W. and T.N. discussed the results and commented on the manuscript. J.Z., G.J. G.W. and T.N. wrote the manuscript with input from other authors. G.J. and P.G. revised the final manuscript. T.N. supervised the entire project.

## Supplementary Figures

**Supplemental Fig 1.**
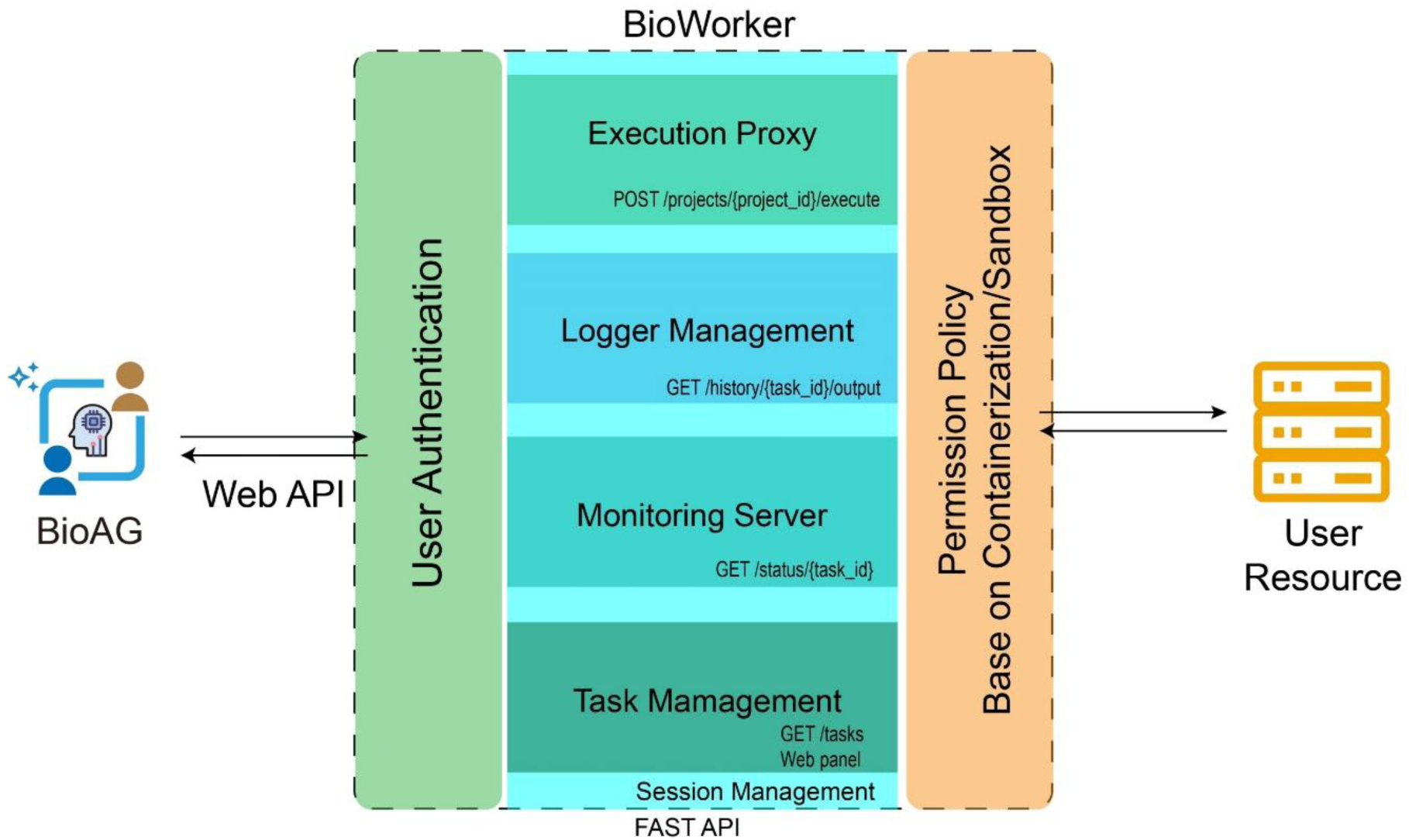
Overview of BioWorker execution proxy. BioWorker is an execution proxy constructed using FastAPI, incorporating an authentication module and session management module to support multi-session task, along with process permission management achieved through containerization or sandbox technologies. BioWorker exposes HTTP APIs for user-side BioAG access or other clients.

**Supplemental Fig 2.**
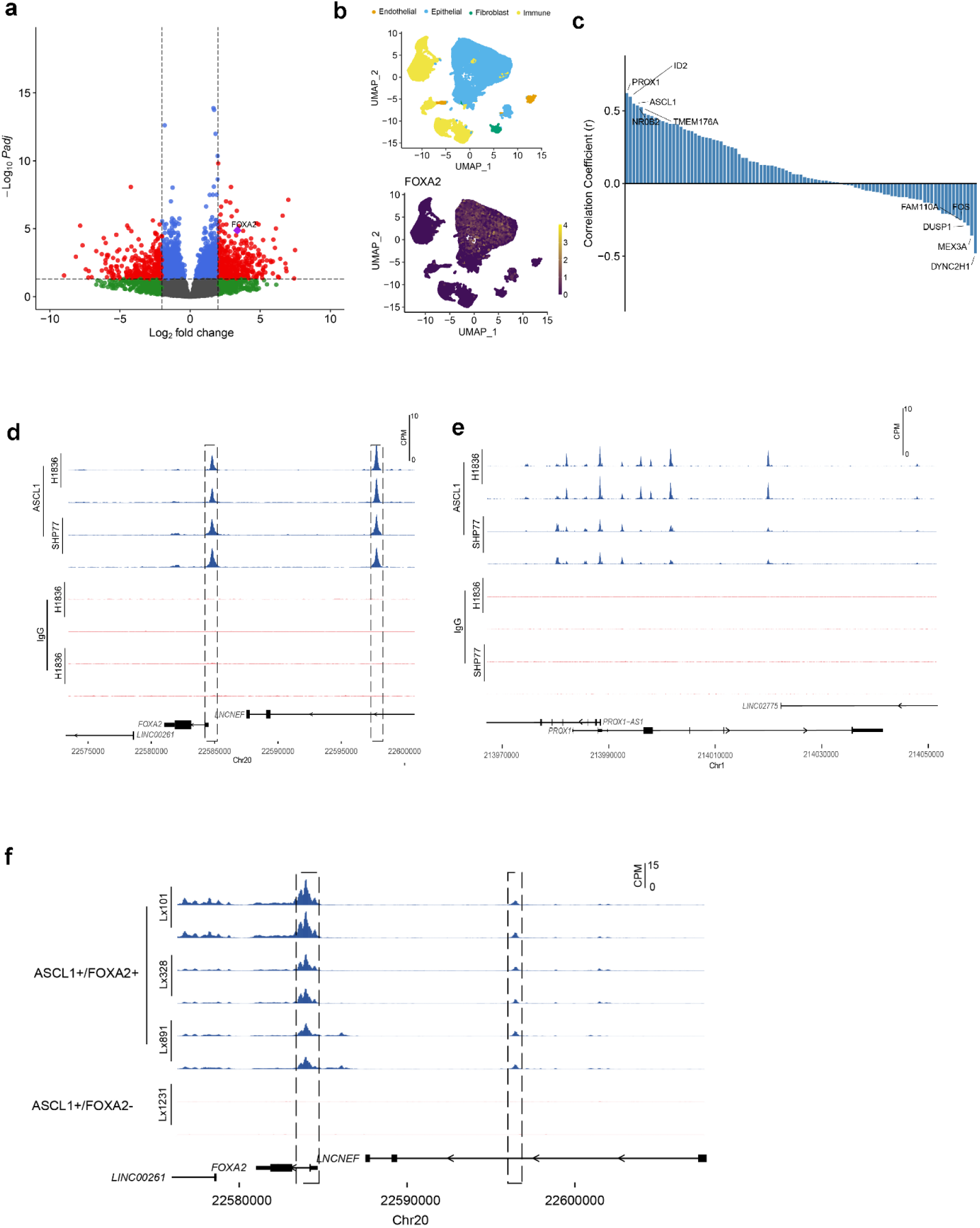
BioGAIP Achieves High-Quality Reproduction of Published Analysis Results. (a) Volcano plot of DEGs in bulk RNA-seq data of met-associated primary SCLC and never-met primary SCLC. (b) UMAP of cell annotation and *FOXA2* expression level in each component based scRNA-seq. (c) Correlation plot of the top 100 highly expressed genes in *FOXA2^+^* vs. *FOXA2^-^* cells from Kawasaki et al. gene set, relative to *FOXA2* expression defined by Kawasaki *et al*^31^. (d,e) Density plot of *ASCL1* ChIP-seq gene loci at the *FOXA2* and *PROX1* gene loci in two SCLC cell lines (H1836 and SHP-77). (f) Density plot at the *FOXA2* locus of ATAC-seq data derived from *ASCL1*^+^ *FOXA2*^+^ PDX vs *ASCL1*^+^ *FOXA2*^-^ PDX tumors.

**Supplemental Fig 3.**
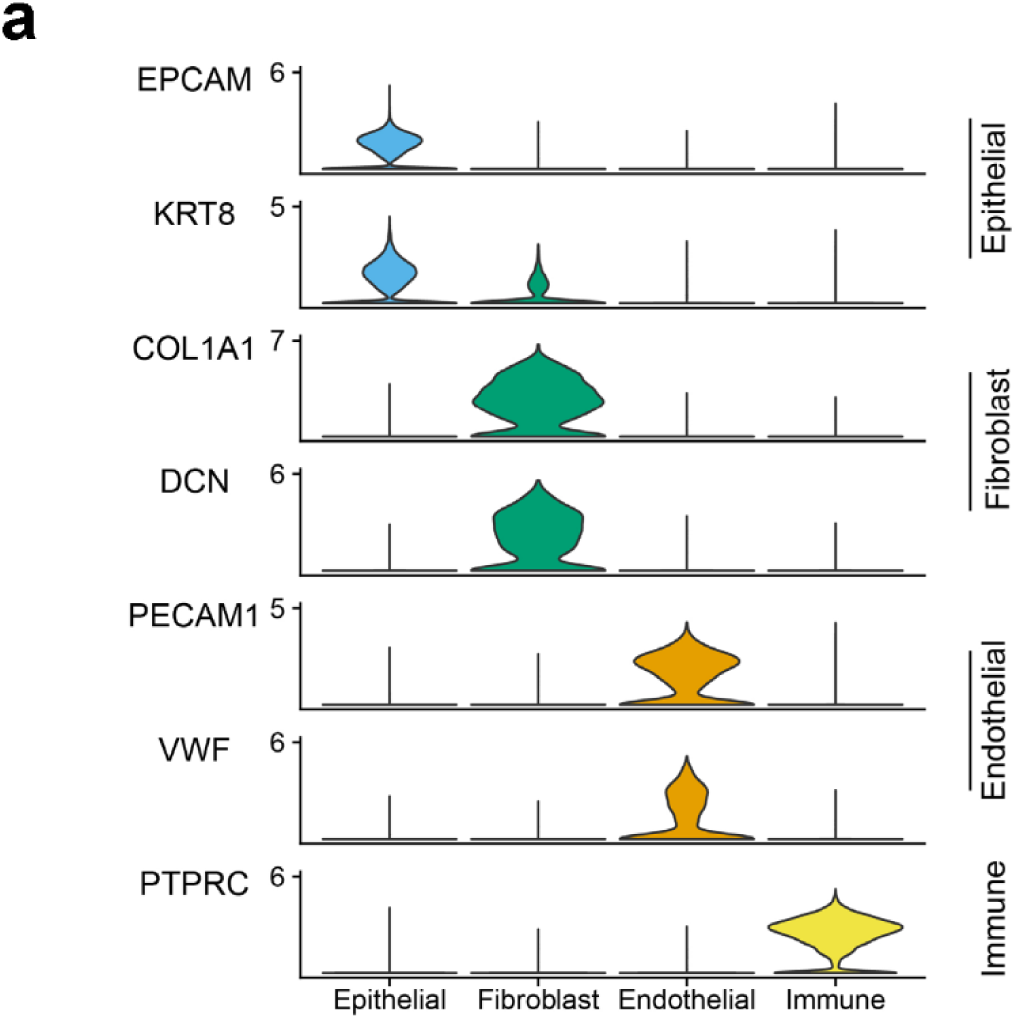
BioGAIP Achieves High-Quality Cell annotation. (a) Violin plots of expression levels for marker genes reported in literatures^31,50^.

**Supplemental Fig 4.**
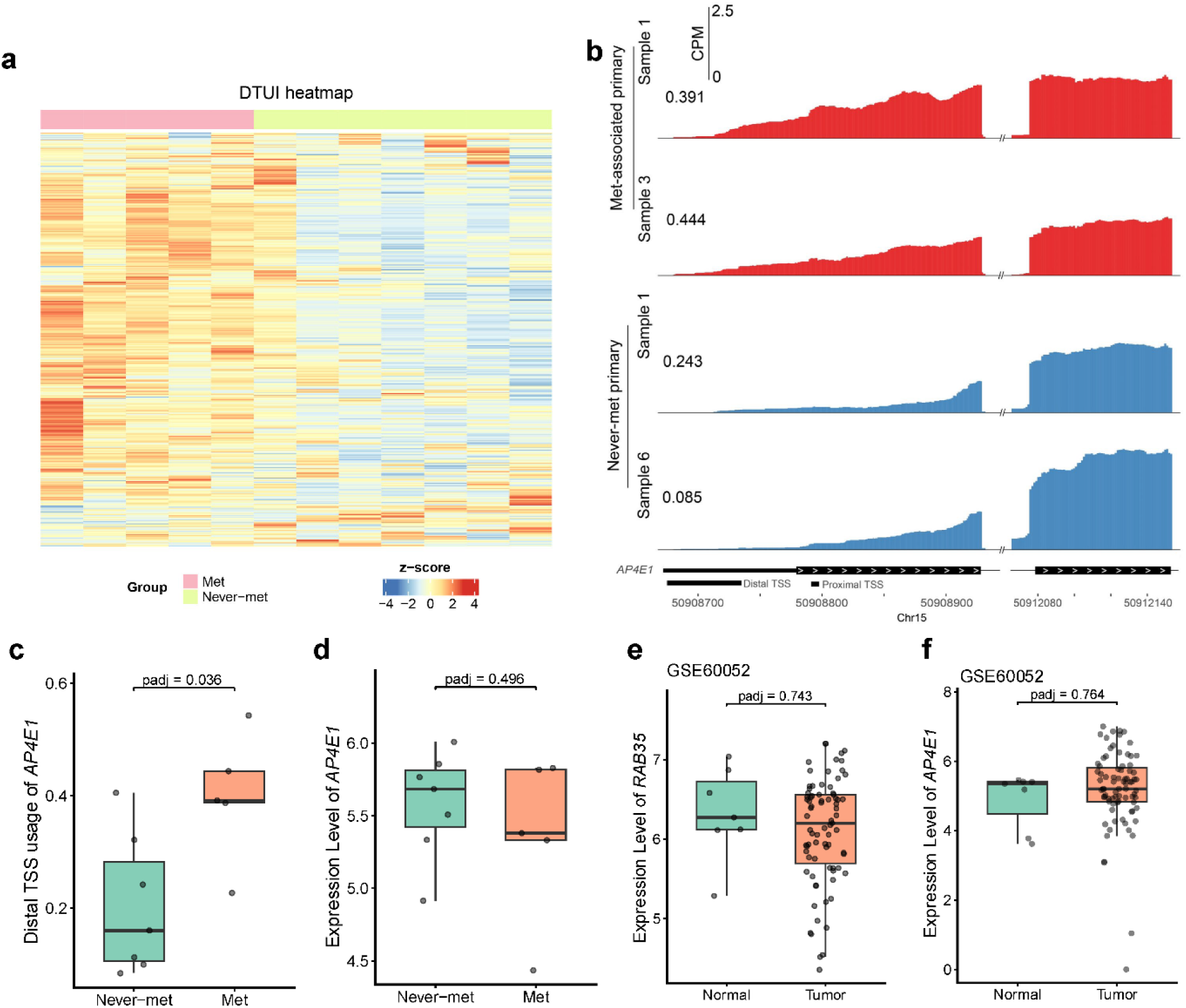
BioGAIP Identifies Dynamic Transcription Initiation Events in SCLC. (a) Heatmap of differential ATTSS events based on DTUI values. (b) RNA-seq density plots show the high distal TSS usage of *AP4E1* in two met-associated primary tumors and two never-met primary tumors. (c) The comparison of distal TSS usage of *AP4E1* between never-met primary and met-associated primary tumor. (d) The comparison of gene expression of *AP4E1* between never-met primary and met-associated primary tumor. (e-f) The comparison of gene expressions of *RAB35* (e) and *AP4E1* (f) between normal and sample tumor in GES60052. Statistical significance of distal TSS usage was assessed using the DATTS^28^, statistical significance of gene expression was assessed using the DESeq2^49^.

