## Supplementary material for "BioGAIP: A Scalable, User-Friendly and Robust LLM-Powered Multi-Agent System for Automated Bioinformatics Tasks": Supplement material 1.pdf

### RNA-seq

#### Prompt:

You are an expert bioinformatician tasked with processing RNA-seq sequencing data from the dataset GSE281525_rna_seq_kawasaki_et_al_2025. The data is stored in the directory <input fastq directory>, which includes RNA-seq data for non-metastatic tumor samples (in the subdirectory Never-met_rna_seq) and metastatic tumor samples (in the subdirectory Met_rna_seq). The human reference genome hg38 and its corresponding GTF annotation file are available in <reference genome directory>.

Perform the following pipeline steps on the data:

Conduct quality control (QC) using appropriate tools such as FastQC or MultiQC.

Align the reads to the hg38 reference genome using STAR.

Quantify transcripts using featureCounts.

Perform differential expression analysis using DESeq2 to identify genes with differential expression between non-metastatic (Never-met_rna_seq) and metastatic (Met_rna_seq) tumor samples.

All outputs must be saved in <output directory>. If you need to create or manage environments (e.g., Conda envs), store them in <env directory>

You have access to <core number> CPU cores for parallel processing where applicable.

Expected Outputs:

Aligned BAM files for each sample.

Transcript quantification results (e.g., count matrices).

A list of differentially expressed genes, including fold changes, p-values, and adjusted p-values.

Constraints:

ONLY ASSIGN ONE TASK EACH TIME!

ONLY THE OUTPUT DIRECTORY AND ENVIRONMENTS DIRECTORY ARE WRITEABLE; do not write to any other paths.

### ChIP-seq

#### Prompt:

The chip-seq sequencing data is stored in the folder <input directory>. This data originates from the H1836 and SHP77 cell lines. The reference genome is located in <reference genome directory>, and blacklists for various reference genomes are available in <chip-seq blacklist directory>.

Perform quality control and analyze the data. Identify peaks in the ASCL1 gene binding regions separately for the H1836 and SHP77 cell lines, and determine the consensus peaks between the two replicates. Technical replicates may be merged as needed.

Save all outputs in <output directory>.

If environments need to be created, store them in <env directory>.

You have total <core number> cores for parallel processing where applicable.

Retain necessary intermediate files. Note that write permissions are limited to the output folder only.

### ATAC-seq

#### Prompt

You are an expert in ATAC-seq data analysis.

The raw ATAC-seq FASTQ files for FOXA2+ and FOXA2- PDX tumor samples are stored in:

<input directory>

File naming convention:

{sample_id}_{sample_name}_{sample_type:FOXA2+/-}_{biologic_rep_id}_tech{tech_rep_id}_{reads_id}.fastq.gz

Example: GSM8622652_x328_FOXA2+_rep1_tech1_1.fastq.gz

Reference genome directory: <reference genome directory>

ATAC-seq blacklist directory: `<atac blacklist directory>`

Please perform the following analysis steps:

1. Quality control of raw reads and align clean reads to the hg38 reference genome.
2. Merge technical replicates into biological replicates.
3. Call consensus peaks between the two biological replicates for each sample.
4. Generate separate consensus peak sets for FOXA2+ samples and FOXA2- samples.
5. Identify differential accessible peaks between FOXA2+ and FOXA2- groups.
6. Keep all necessary intermediate files.

All output files must be saved in:

`<output directory> `

If a conda environment is needed, create and save it in:

`<env directory> `

You are only allowed to write to the above output directory and environment directory.

You have total <core number> CPU cores to use, please try to fully use all CPU resource.

### scRNA-seq

#### Prompt 1: scRNA-seq Upstream analysis

Your task is process the provided scRNA-seq data from small cell lung cancer (SCLC) patients.

Data Details:

Input data path: <input directory>

File naming convention: {access_id}_{sample_id}_{tissue_type}_{sex}_age_{age}_{rep_type}_{reads_id}.fastq.gz

Files may include technical replicates.

Available reference genomes: <reference genome> and <refdata-gex-GRCh38-2024-A reference genome path>.

Cell Ranger 8.0.1.tar.gz has been download to `<cell ranger tar.gz path>`

Please carefully check mapping for sra dumped fastq file and cellrange input read 1, read 2 and Index 1.

(IMPORTANT: _2.fastq.gz is R1, _3.fastq.gz is R2, _1.fastq.gz is I1 in this dataset)

It is recommended that you link all the files to a new folder and modify the linked file names to a format compatible with cellranger

Output Directory:

Save all results to: <output directory>

Environment Setup (if needed):

Create environment in: <env directory>

Requirements:

1. Alignment with CellRanger:

Use CellRanger to align all data, retain BAM files, and generate expression matrices.

2. Import and Merge in Seurat:

Use Seurat to import all expression matrices, merge them, add necessary annotations (meta data), and handle batch effects.

You have total <core number> cores for parallel processing where applicable. Please try to fully use that.

Retain necessary intermediate files. Note that write permissions are limited to the output folder only.

### Prompt 2: scRNA-seq Downstream analysis

The following RDS file contains single-cell RNA-seq data from small cell lung cancer patient samples. The data has undergone necessary preprocessing, such as clustering and dimensionality reduction, and is saved as a Seurat object.

RDS file path: <input Seurat RDS file>

Complete the following tasks step by step, using only the actual data loaded from the RDS file. Do not assume, invent, or hallucinate any data, results, or annotations not directly supported by the loaded data and your verified knowledge of marker genes.

Annotate the cells by assigning them to one of the following categories: Epithelial, Immune, Endothelial, or Fibroblast. To do this, first identify marker genes for each cluster using standard Seurat methods (e.g., FindAllMarkers), then apply your knowledge of established cell-type-specific markers to assign annotations accurately.

Generate a violin plot illustrating the expression level differences of the FOXA2 gene across the annotated Epithelial, Immune, Endothelial, and Fibroblast cell types.

Subset the data to include only cells annotated as Epithelial. Classify these cells into FOXA2 high-expression (FOXA2+) and FOXA2 low-expression (FOXA2-) groups based on FOXA2 expression levels (e.g., using a threshold derived from the data distribution). Identify the differentially expressed genes between these two groups using standard differential expression analysis (e.g., FindMarkers in Seurat).

Please install presto package to accelerate FindMarkers function

All cells must be assign to Immune,Epithelial,Fibroblast or Endothelial

Save all outputs—including any normalized tables, differential expression results tables, ranked lists of significant genes (with p-values and fold changes), and the violin plot as PNG or PDF—to the directory:

<output directory>

You have write permissions only in the specified output directories. If creating virtual environments is necessary, place them in:

<env directory>

Display key code snippets and intermediate results in your response for verification, but ensure all final files are written exclusively to the designated output path. Base all analyses and outputs strictly on the loaded data without extrapolation.

**Note, this cell annotation tasks in the downstream analysis pipeline (Prompt 2) requires manual curation of BioGAIP-generated cell annotations followed by iterative refinement through direct interaction with the agents. Because these human–agent interactions exert a major influence on the final annotations and because it is inherently difficult to apply an objective, binary classification of individual cell annotations as successes or failures, the Prompt 2 results were excluded from Table 2. Downstream analysis results generated with Qwen-max were used to produce Supplementary Figs. 2b and 3a.**

### WGS

#### Prompt:

You are an experienced bioinformatician. Now process the WGS sequencing data from cell lines according to the following requirements. Strictly follow these steps exactly; do not add, omit, or assume any extra steps, tools, or paths.

1. Perform necessary quality control and preprocessing on the sequencing data.

2. Align the sequencing data to the reference genome using bwa.

3. Perform quality control and processing on the mapped BAM files (e.g., duplicate removal).

4. Use GATK to perform variant analysis for each sample and merge all analysis results.

5. Perform quality control on the variant analysis results.

6. Retain all necessary intermediate results and logs.

Input sequencing data location: <input directory>

Reference genome location: <reference fasta>

Output directory: <output directory>

Conda environment: <env directory> (use only if needed; prefer pre-installed packages).

You have total 29 cpu cores to use

Note: You only have write permissions to the output folder and the conda folder.

### IPA analysis

### Prompt

You are an experienced bioinformatics analyst specializing in RNA-seq data processing and alternative polyadenylation analysis.

Background

Alternative polyadenylation (APA) is a post-transcriptional regulatory mechanism occurring at the 3' end of mRNA. Intronic polyadenylation (IPA) is a subtype of APA that typically alters the coding region of mRNA. Quantitative methods for IPA have been published in the following article:

Zhao Z, Xu Q, Wei R, Wang W, Ding D, Yang Y, Yao J, Zhang L, Hu YQ, Wei G, Ni T. Cancer-associated dynamics and potential regulators of intronic polyadenylation revealed by IPAFinder using standard RNA-seq data. Genome Res. 2021 Sep 2. doi: 10.1101/gr.271627.120. PMID: 34475268.

The associated tool, IPAFinder, is open-sourced at: https://github.com/ZhaozzReal/IPAFinder

Small cell lung cancer (SCLC) is a disease with poor prognosis and limited treatment options. SCLC is highly metastatic, but the underlying mechanisms are unclear. We have collected RNA-seq data from metastatic (Met) SCLC primary tumors and never-metastatic (Never-met) tumors, aligned to the human reference genome hg38. The aligned BAM files are located in: <input bam path>

Task Instructions

1. Visit the IPAFinder GitHub repository to retrieve the software's installation and usage guidelines. Focus specifically on the methods for "Detect IPA sites and quantify their usages" and "Infer statistically differential usage of IPA sites."

2. Strictly adhere to the documented steps from the repository without assuming or inventing any undocumented features or parameters. If any information is unclear, report it precisely without speculation.

Requirements:

1. Install and configure the IPAFinder software. If environment management (e.g., Conda) is needed, create and store environments in <env directory>.

2. Use IPAFinder to detect and quantify IPA sites and their usages for each sample in the

provided BAM files.

Identify and list statistically differential IPA events between the Met and Never-met groups.

Retain all necessary intermediate files for reproducibility.

Utilize up to <core number> CPU cores for parallel processing where supported by the tool.

Save all outputs in <output directory>.

Expected Outputs

1. Quantification results for IPA events across samples.

2. List of differential IPA events between Met and Never-met groups.

Respond step-by-step, documenting each action taken, commands executed, and any errors encountered. Base all actions solely on the provided background, repository documentation, and data paths, do not introduce external assumptions or hallucinations.

### ATTSS analysis

#### Prompt

Your goal is to analyze RNA-seq data for alternative tandem TSS events in small cell lung cancer (SCLC) samples using the DATTSS tool, strictly following the provided instructions and sources without adding or inferring any extra details. Base all actions on verified information from the specified GitHub repository and avoid hallucinations by using tools to fetch and verify data.

Background:

Alternative tandem transcription initiation (Dynamic analyses of Alternative Tandem TSS events) is an important transcriptional regulatory mechanism. Related quantitative methods have been published in the following article:

Zhao Z, Chen Y, Zou X, Lin L, Zhou X, Cheng X, Yang G, Xu Q, Gong L, Li L, Ni T. Pan-cancer transcriptome analysis reveals widespread regulation through alternative tandem transcription initiation. Sci Adv. 2024 Jul 12;10(28):eadl5606. PMID: 38985880.

The related tool (DATTSS) is open-sourced at: https://github.com/ZhaozzReal/DATTSS

SCLC is a disease with poor prognosis and limited treatment options. SCLC is highly prone to metastasis, and the underlying mechanisms are unclear. We have collected RNA-seq data from metastatic (Met) SCLC primary tumors and never-metastatic (Never-met) tumors, aligned to the human reference genome hg38. The aligned BAM files are located at: <input bam file>

First, access the DATTSS GitHub repository and extract the installation and usage instructions from its readme.md, particularly for the method to compare alternative tandem TSS usage between conditions. Do not assume or invent any details not present in the repository.

Requirements:

Install and configure the DATTSS software in new environment, following the instructions fetched from GitHub.

Use the DATTSS software to quantify alternative tandem TSS events in each sample.

Identify the list of alternative tandem TSS events that differ between Met and Never-met groups.

Retain necessary intermediate files.

All outputs must be saved in <output directory>. If you need to create or manage environments (e.g., Conda envs), store them in <env directory>.

You have access to <core number> CPU cores for parallel processing where applicable.

Constraints:

ONLY ASSIGN ONE TASK EACH TIME!

ONLY THE OUTPUT DIRECTORY AND ENVIRONMENTS DIRECTORY ARE WRITEABLE; do not write to any other paths.

### GEO query

#### Prompt

You are a bioinformatics expert tasked with analyzing the GEO public dataset GSE60052, which contains RNA-seq data from 79 small cell lung cancer (SCLC) samples and 7 normal control samples. SCLC is a tumor with complex pathogenesis and poor prognosis. Follow these steps precisely, using verified tools and data sources to avoid hallucinations or assumptions. Base all actions on factual data retrieved from reliable databases, and document your reasoning step by step.

1. Access the GEO database via appropriate tools to retrieve and summarize relevant information for the accession number GSE60052, including dataset description. Do not assume any details; verify everything from the source.

2. Directly Download the gene expression data from the GEO database web page if needed (use wget or other tools). Analyze its content and structure, including file format, sample groups, and any preprocessing notes. Output a clear summary of the findings.

3. Use the limma R package to perform differential expression analysis. Identify differentially expressed genes between the tumor samples (SCLC) and the paired/control samples (normal). Apply standard statistical methods: normalize data, fit models, and use adjusted p-values (e.g., FDR < 0.05) for significance. Explain your code and results step by step, without fabricating data.

Save all outputs, including files, logs, and results, in <output directory>. If creating or managing environments (e.g., Conda envs), store them in <env directory>.

Proceed step by step, and only use tools or code that directly support the tasks. If any information is unclear or inaccessible, note it explicitly without guessing.

### Cell Scenescence Evaluation

#### Prompt

You are an experienced bioinformatician tasked with evaluating cellular senescence using the human universal senescence index (hUSI) on public RNA-seq data.

Background: Cellular senescence is involved in various physiological and pathological processes, including aging and cancer. Multiple methods exist for assessing cellular senescence, such as the human universal senescence index (hUSI, Publication: Wang, J., Zhou, X., Yu, P. et al. A transcriptome-based human universal senescence index (hUSI) robustly predicts cellular senescence under various conditions. Nat Aging (2025).). The related code is open-sourced at: https://github.com/WJPina/HUSI. The GEO database's public dataset GSE130727 contains RNA-seq sequencing results for 8 different cellular senescence models.

Complete the following tasks step by step:

Download the Supplementary file for GSE130727 from the GEO database, decompress it, and integrate the data according to the meta information provided on the webpage.

Read the hUSI literature and GitHub documentation, then download and configure hUSI.

Use hUSI to evaluate the cellular senescence index for the different senescence models in the GSE130727 data.

Retain intermediate results from the hUSI execution process.

Paths:

Output directory: <output directory>

Conda environment: <env directory>.

**Note: It is possible that, due to some network issues encountered during the testing, the following content was added to certain test prompts:**

Considering that the internet connection might be unstable, please try to download GSE130727_RAW.tar several times if error.
