## Supplementary material for "BioGAIP: A Scalable, User-Friendly and Robust LLM-Powered Multi-Agent System for Automated Bioinformatics Tasks": Supplemental Table 1.pdf

### Supplemental Table 1. BioGAIP Compatibility Test Results of LLM Models and Platforms

| Family | Model | Compatibility (Windows 11*) (Without Containerization) | Compatibility (Windows 11 WSL2) (Containerization) | Compatibility (Ubuntu 20.04) (Containerization) |
|---|---|---|---|---|
| Grok | grok-4-fast-reasoning | Mostly Functional | Fully | Fully |
| | grok-4-0709 | Mostly Functional | Fully | Fully |
| | grok-code-fast-1 (For coding) | Fully | Fully | Fully |
| Gemini | gemini-2.5-flash-preview-09-2025 | Mostly Functional | Mostly Functional | Mostly Functional |
| | gemini-3-flash-preview | Mostly Functional | Mostly Functional | Mostly Functional |
| QWen | qwen-plus-2025-12-01 | Fully | Fully | Fully |
| | qwen3-max-2025-09-23 | Fully | Fully | Fully |
| DeepSeek | deepseek-chat (V3.2-Exp) | Fully | Fully | Fully |
| | deepseek-reasoner (V3.2-Exp) | Mostly Functional | Fully | Fully |
| ChatGPT | gpt-4.1-mini | Fully | Fully | Fully |
| | gpt-4o | Fully | Fully | Fully |

* On the Windows platform, BioWorker cannot run in non containerization support, whereas the other components remain functional. The compatibility tests presented in the table for this scenario include only the operational components (BioAG and BioLauncher).
