## Supplementary material for "BioGAIP: A Scalable, User-Friendly and Robust LLM-Powered Multi-Agent System for Automated Bioinformatics Tasks": Supplemental Table 2.pdf

| The Public Data Sources Used in the Article | | |
|---|---|---|
| Task name | Data accession | Ref |
| ATAC-seq | GSE281523 | Kawasaki et. al., 2025 |
| ChIP-seq | GSE281524 | Kawasaki et. al., 2025 |
| RNA-seq | GSE281525 | Kawasaki et. al., 2025 |
| scRNA-seq | GSE281740 | Kawasaki et. al., 2025 |
| WGS | SRR8788981,SRR8670685,SRR8652105 | Barretina J et al., 2012;Ghand et.al., 2019 |
| IPA | GSE281525 | Kawasaki et. al., 2025 |
| ATTSS | GSE281525 | Kawasaki et. al., 2025 |
| GEO Query | GSE60052 | Jiang et. al., 2016 |
| Cell Senescence Evaluation | GSE130727 | Casella et. al., 2019 |
