## Supplementary material for "BioGAIP: A Scalable, User-Friendly and Robust LLM-Powered Multi-Agent System for Automated Bioinformatics Tasks": Supplementary Data Detail.pdf

Table of supplementary content

- Supplemental table 1: BioGAIP Compatibility Test Results of LLM and Platforms
- Supplemental table 2: The Public Data Sources Used in the Article

Materials of supplementary content

- Supplemental material 1: The List of Prompts Used in This Study

Videos of supplementary content

- Supplemental video 1: BioLauncher: Comprehensive Graphical Deployment Support for BioGAIP
- Supplemental video 2: BioGAIP Completes RNA-seq Analysis Tasks with Minimal Human Intervention
